# Tenofovir Activation is Diminished in the Brain and Liver of Creatine Kinase Brain-Type Knockout Mice

**DOI:** 10.1101/2023.09.25.559370

**Authors:** Colten D. Eberhard, Eric P. Mosher, Namandjé N. Bumpus, Benjamin C. Orsburn

## Abstract

Tenofovir (TFV) is a nucleotide reverse transcriptase inhibitor prescribed for the treatment and prevention of human immunodeficiency virus infection, and the treatment of chronic hepatitis B virus infection. Here, we demonstrate that creatine kinase brain-type (CKB) can form tenofovir-diphosphate (TFV-DP), the pharmacologically active metabolite, in vitro, and identify nine missense mutations (C74S, R96P, S128R, R132H, R172P, R236Q, C283S, R292Q, and H296R) that diminish this activity. Additional characterization of these mutations reveal that five (R96P, R132H, R236Q, C283S, and R292Q) have ATP dephosphorylation catalytic efficiencies less than 20% of wild-type (WT), and seven (C74S, R96P, R132H, R172P, R236Q, C283S, and H296P) induce thermal instabilities. To determine the extent CKB contributes to TFV activation in vivo, we generated a CKB knockout mouse strain, *Ckb^tm1Nnb^*. Using an in vitro assay, we show that brain lysates of *Ckb^tm1Nnb^* male and female mice form 70.5% and 77.4% less TFV-DP than wild-type brain lysates of the same sex, respectively. Additionally, we observe that *Ckb^tm1Nnb^* male mice treated with tenofovir disoproxil fumarate for 14 days exhibit a 22.8% reduction in TFV activation in liver compared to wild-type male mice. Lastly, we utilize mass spectrometry-based proteomics to elucidate the impact of the knockout on the abundance of nucleotide and small molecule kinases in the brain and liver, adding to our understanding of how loss of CKB may be impacting tenofovir activation in these tissues. Together, our data suggest that disruptions in CKB may lower levels of active drug in brain and liver.

**ABSTRACT GRAPHIC:** 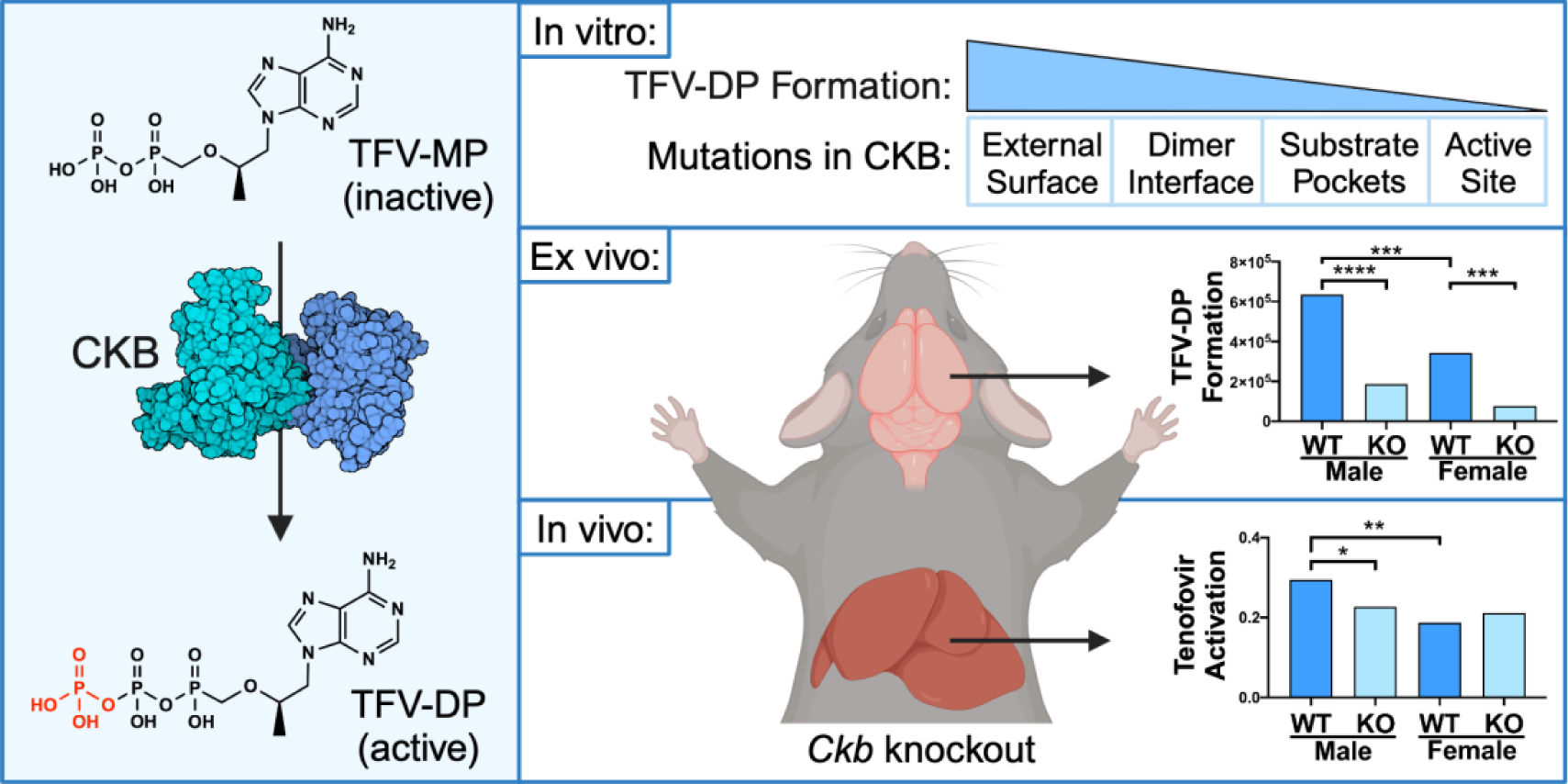

## INTRODUCTION

Nucleot(s)ide analogues are a large class of antiviral drugs that are prescribed to treat or prevent many viral infections. Tenofovir (TFV) is an acyclic nucleoside phosphonate analogue of adenosine 5’-monophosphate that is sequentially phosphorylated to inhibit human immunodeficiency virus (HIV) reverse transcriptase and hepatitis B virus (HBV) polymerase.^1,2^ TFV is formulated as tenofovir disoproxil fumarate (TDF) and tenofovir alafenamide, which are prodrugs approved as monotherapies to treat HIV and chronic HBV infections, as well as components of combination therapies for HIV pre-exposure prophylaxis (PrEP).^1,3,4^ Both prodrugs must undergo hydrolysis, either by cathepsin A or carboxylesterase 1, to become TFV, which is then phosphorylated twice to become the pharmacologically active metabolite, tenofovir-diphosphate (TFV-DP).^5,6^ Once incorporated into nascent viral DNA, TFV-DP terminates the elongation step of viral DNA synthesis.^2,7^

While the efficacy of TFV-based PrEP remains high, HIV breakthrough infections can still occur and are often attributed to a lack of adherence.^8–12^ Recent studies have begun correlating the concentration of TFV-DP in dried blood spots to the number of doses taken each week, an unbiased way of measuring adherence.^13^ However, a 2022 study investigating PrEP efficacy in adolescent girls and young women reported that three HIV seroconverters had TFV-DP levels in dried blood spots equating to the individual taking 4-6 doses per week, an amount that reduces risk of infection 96%.^10,14^ Similarly, two cases have been reported where an individual taking tenofovir-based PrEP acquired tenofovir-susceptible HIV infection despite maintaining high adherence to the regimen, also confirmed through dried blood spot analyses.^15,16^ These data bring forth a disconnect between adherence or TFV-DP levels in blood and overall efficacy.

Adenylate kinase 2 (AK2) is the primary enzyme that phosphorylates TFV to the intermediate metabolite tenofovir-monophosphate (TFV-MP).^17^ In peripheral blood mononuclear cells and vaginal tissue, pyruvate kinase isoenzymes PKLR and PKM catalyze TFV-DP formation, whereas in colon tissue, creatine kinase muscle-type (CKM) is principally responsible.^17^ Since the enzymes contributing to the formation of TFV-DP are tissue-dependent, we postulate that interindividual variability in these TFV activating enzymes may cause asymmetric concentrations of TFV-DP between systemic circulation and peripheral tissues, reducing tissue-specific efficacy.^18^ Previous in vitro studies have elucidated the impact of naturally occurring mutations in AK2 and CKM on the in vitro formation of TFV-MP and TFV-DP, respectively.^19,20^ To expand our understanding of tissue-specific protection and TFV efficacy, it is important to identify TFV activating enzymes in tissues susceptible to HIV and HBV infection.

We hypothesize that creatine kinase brain-type (CKB) may play a role in TFV activation as it shares 80% amino acid sequence identity to CKM. In humans, the *CKB* gene is expressed in tissues relevant to TFV disposition, including liver, vagina, and colon.^21^ Of particular interest, the abundance of CKB protein is high in astrocytes, a cell type that harbors HIV infection in the brain.^22–24^ Like other creatine kinase enzymes, CKB regulates cellular energy homeostasis by buffering ATP concentrations. In subcellular regions where energy is being rapidly consumed (e.g., surrounding ATPases), CKB transfers the phosphate group from phosphocreatine to ADP, increasing local concentrations of ATP.^25^ The role of CKB in the creatine/phosphocreatine cycle has been studied in various tissues and dysregulation of CKB via mutations, oxidation, or abundance alterations are associated with several disease states including Alzheimer’s and Huntingtin’s diseases, and cancer metastases.^24,26–34^ However, the contribution of CKB to xenobiotic biotransformation remains underexplored, as most studies have only investigated the contribution of CKM.^35–38^

In this work, we investigate the contribution of CKB in TFV activation and characterize how naturally occurring mutations disrupt the enzymatic function and stability, in vitro. We establish a CRISPR/Cas9-mediated CKB knockout mouse strain, *Ckb^tm1Nnb^*, to profile the tissue-specific pharmacological role of CKB. Using mass spectrometry-based proteomic analyses, we compile a list of nucleotide and small molecule kinases found in brain and liver and investigate those that are differentially abundant to better understand how loss of CKB impacts other TFV regulating enzymes. Together, this work implicates a role for CKB in TFV activation and emphasizes how alterations in the function of CKB could directly or indirectly reduce TFV activation, potentially effecting TFV efficacy.

## RESULTS AND DISCUSSION

### CKB Catalyzes the Formation of TFV-DP in vitro

To determine whether CKB can catalyze the formation of TFV-DP through the transfer of the phosphate group from phosphocreatine to TFV-MP, we employed an in vitro assay using recombinantly expressed and purified CKM, CKB, and CKMT1 proteins. Each enzyme was incubated in the presence of phosphocreatine and TFV-MP and the resulting tenofovir metabolites were detected using ultra-high performance liquid chromatography tandem mass spectrometry (uHPLC-MS). Analysis of the area under the peak curve revealed no significant differences in TFV-DP levels between the CKM, CKB, or CKMT1 catalyzed reactions **(Figure 1A)**. To test for intrinsic enzymatic differences in the canonical reaction, the proteins were incubated with phosphocreatine and ADP and ATP was detected by uHPLC-MS. Results indicate there were no significant differences in the formation of ATP between isoenzymes **(Figure 1B)**. Overall, we observed that CKB can catalyze the formation of ATP and TFV-DP in vitro to a similar extent as CKM, an enzyme known to contribute to TFV-DP formation in colon tissue ex vivo.^17^

**Figure 1.**
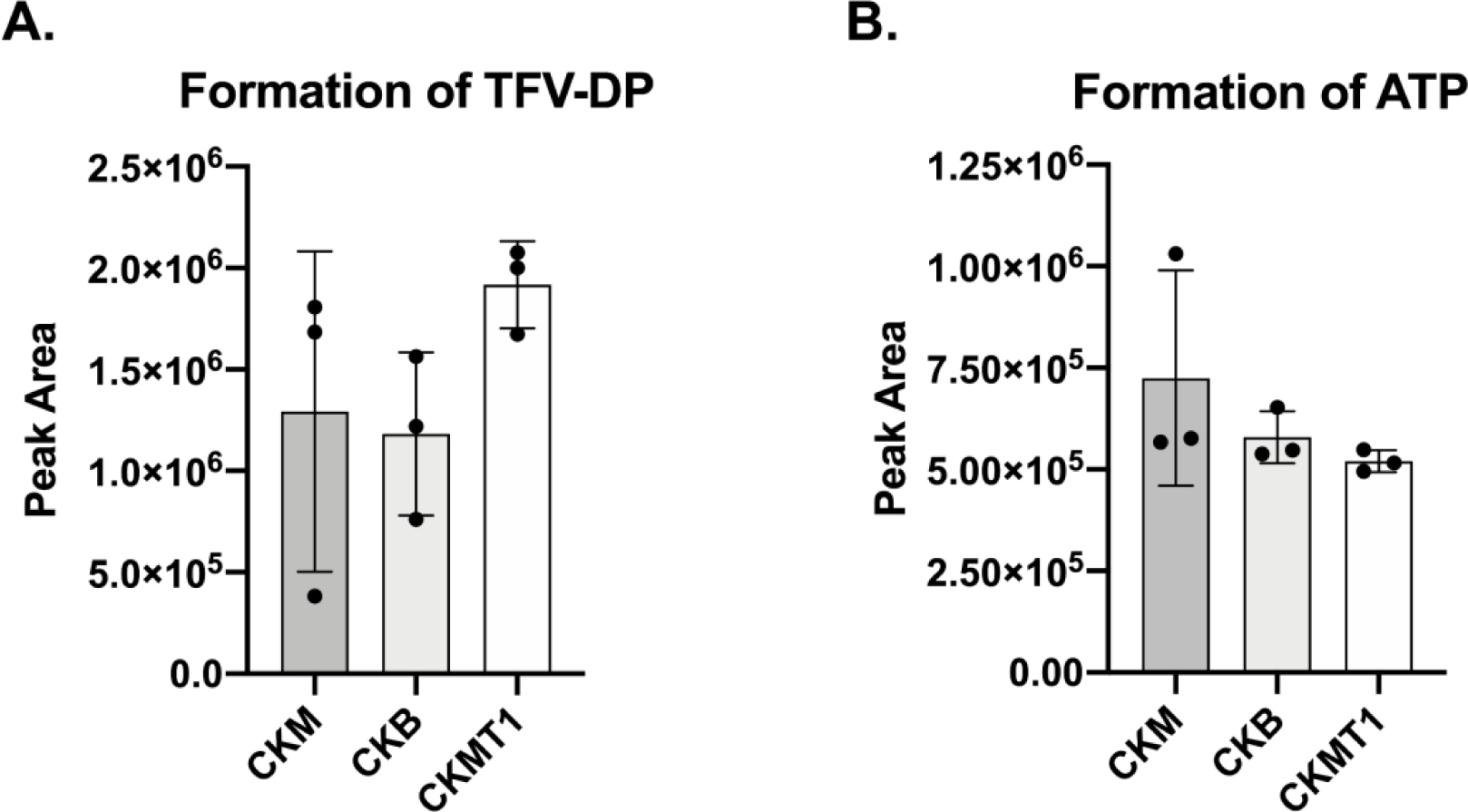
Cytosolic (CKM and CKB) and mitochondrial (CKMT1) creatine kinases show similar activities in the formation of (**A**) TFV-DP and (**B**) ATP. Recombinantly expressed and purified enzymes were incubated with phosphocreatine and TFV-MP or ADP. Metabolites were detected using uHPLC-MS and area under the peak curve was used to measure metabolite levels, (n=3). Error bars represent standard deviation. Statistical analyses were performed using a one-way Brown-Forsythe and Welch ANOVA with Dunnett’s multiple comparisons test. Despite trends, no comparisons were found to be significantly different.

### Mutations in CKB Diminish Enzymatic Activity in vitro

To understand the impact of genetic variation on CKB activity, fifteen naturally occurring mutations found in the human *CKB* gene were selected for in vitro characterization (**Supplemental Table 2**). C283S was included in these analyses as a negative control, as it has been previously determined to have no enzymatic activity.^39^

Mutant CKB enzymes were incubated in the presence of phosphocreatine and TFV-MP or ADP. Reaction products, ATP or TFV-DP, were detected by uHPLC-MS and levels were compared to WT CKB. Mutations at residues where phosphoryl transfer occurs (R132H, R236Q, and R292Q) and C283S significantly reduced the formation of ATP with average activities of 2.2%, 2.5%, 30.8%, and 5.5% of WT, respectively **(Figure 2A)**. Eight naturally occurring mutations (C74S, R96P, S128R, R132H, R172P, R236Q, R292Q, and H296R) and C283S demonstrated a statistically significant reduction in the formation of TFV-DP with average activities of 20.1%, 6.8%, 30.4%, 1.5%, 21.3%, 1.4%, 2.5%, 5.6%, and 1.4% of WT, respectively **(Figure 2B)**. Together, these data indicate C74S, R96P, S128R, R172P, and H296R exhibited a substrate-specific impact on activity. A similar phenomenon has been noted for mutations in CKM, where it has been proposed that individuals possessing substrate-dependent mutations may not display a clinical phenotype associated with disruptions in canonical pathways, but may be unable to form TFV-DP as readily.^19^ In the solved crystal structure (PDB: 3B6R), Cys74 sits at the bottom of the creatine binding pocket, but does not interact directly with another residue or substrate, while Arg96 interacts indirectly with creatine through a water molecule, and Ser128 and His296 have direct interactions with the adenine ring of ADP.^40^ We suspect the substrate-specific characteristic of these mutations occurs via conformational changes of the binding region or the loss of indirect or direct interactions with certain substrates. Further, we note that disruptions in activity aren’t just substrate- and site-specific, but are mutation-specific. For example, R172P reduced TFV-DP formation 78.7% and R172G only reduced activity 27.2%, while neither mutation had an impact on ATP formation. This is similarly observed with His296 mutations. Thus, predicting the impact of a mutation will require additional work-up to determine clinical relevance.

**Figure 2.**
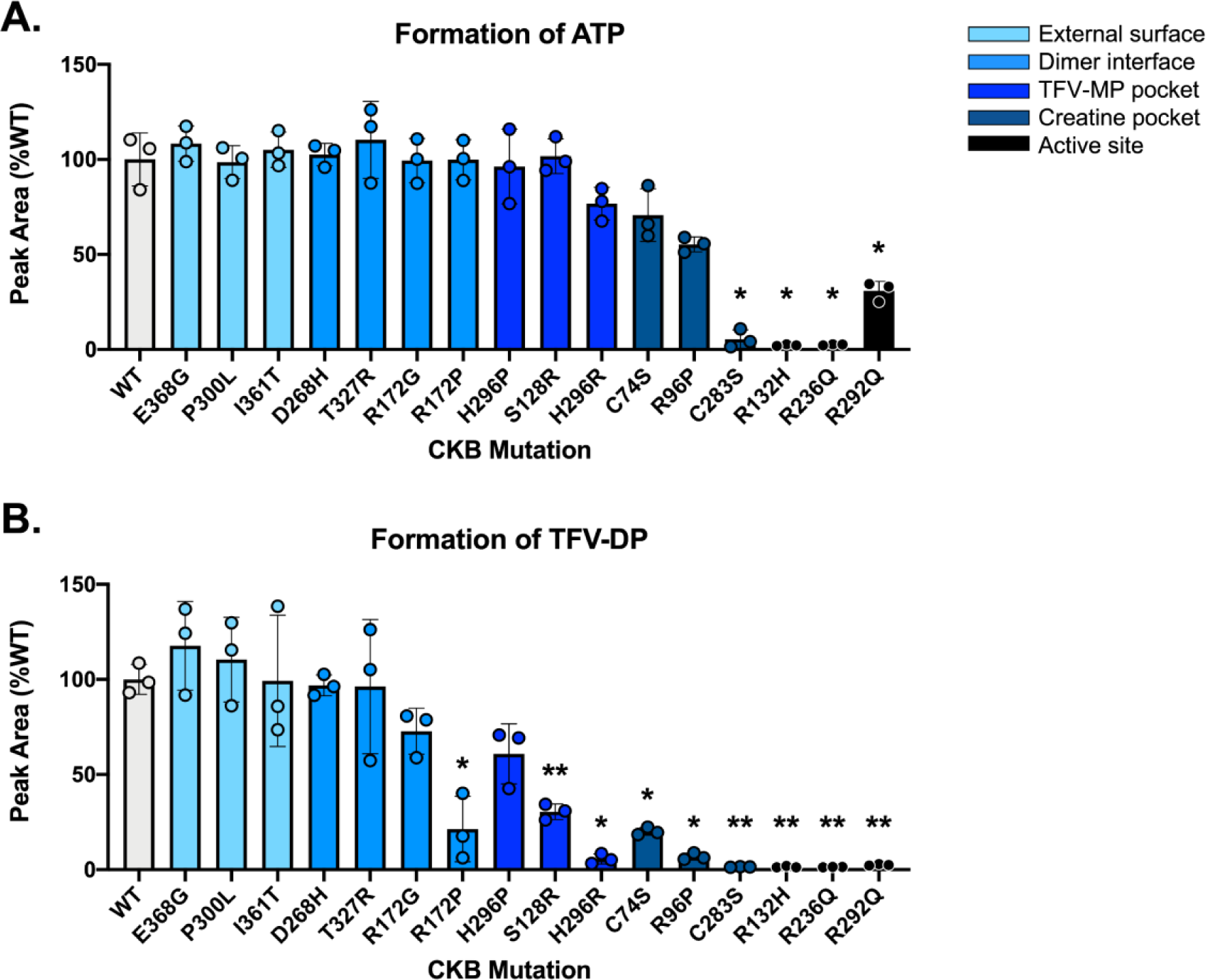
In vitro activity of sixteen recombinantly expressed mutant CKB enzymes in the formation of (**A**) ATP and (**B**) TFV-DP. Phosphocreatine and reaction substrate (ADP or TFV-MP) were incubated with purified enzyme, and ATP or TFV-DP was detected using uHPLC-MS. Metabolite formation is shown normalized to the average wild-type area under the peak curve. Assays were conducted in triplicate and error bars represent standard deviation. Statistical analyses were performed using a one-way Brown-Forsythe and Welch ANOVA with Dunnett’s multiple comparisons test; P-values < 0.05*, 0.01**.

Overall, mutations at active site arginine residues (R132H, R236Q, and R292Q) had the most severe effect on activity, reducing TFV-DP formation 97.5-98.6% compared to WT. Mutations within the phosphocreatine binding pocket (C74S and R96P) and predicted TFV-MP binding pocket (S128R and H296R) also significantly diminished TFV-DP formation (69.6-94.4%). Of note, R172P was the only mutation located in the dimer interface that significantly reduced the formation of TFV-DP.

To characterize how these mutations disrupt enzymatic activity, the reverse canonical enzymatic reaction (ATP dephosphorylation) was exploited in an enzyme coupled system, from which Michaelis-Menten curves were constructed (**Supplemental Figure 1)** and resulting kinetic parameters were extracted **(Table 1)**. Five naturally occurring mutations (R96P, R132H, R236Q, R292Q, and H296R) and C283S exhibited catalytic efficiencies (kcat/Km) less than 20% of WT. The decrease in catalytic efficiency was generally due to increases in Km, signifying these mutations impact substrate binding.

**Table 1.** Michaelis-Menten Kinetic Parameters of Mutant CKB Enzymes.^1^.

| CKB Mutant | $k_{cat}$ ($s^{-1}$) | $K_m$ (mM) | $k_{cat}/K_m$ ($M^{-1}s^{-1}$) | % WT $k_{cat}/K_m$ ($M^{-1}s^{-1}$) |
| --- | --- | --- | --- | --- |
| WT | $15.5 \pm 0.7$ | $2.05 \pm 0.4$ | $7550 \pm 1560$ | 100 |
| C74S<sup>†</sup> | $15.4 \pm 1.2$ | $1.23 \pm 0.5$ | $12500 \pm 5040$ | 166 |
| R96P<sup>†</sup> | $12.1 \pm 1.1$ | $9.11 \pm 2.5$ | $1340 \pm 381$ | 17.6 |
| S128R<sup>†</sup> | $12.0 \pm 0.8$ | $4.16 \pm 1.0$ | $2890 \pm 696$ | 38.3 |
| R132H<sup>†</sup> | N/A | N/A | $159 \pm 12$ | 2.1 |
| R172G | $14.2 \pm 0.5$ | $1.71 \pm 0.3$ | $8350 \pm 1460$ | 111 |
| R172P<sup>†</sup> | $17.2 \pm 1.0$ | $1.77 \pm 0.5$ | $9760 \pm 2670$ | 129 |
| R236Q<sup>†</sup> | $13.7 \pm 1.5$ | $14.7 \pm 4.2$ | $927 \pm 281$ | 12.3 |
| D268H | $17.3 \pm 0.9$ | $1.13 \pm 0.3$ | $15300 \pm 4080$ | 203 |
| C283S<sup>†</sup> | $13.1 \pm 1.3$ | $17.2 \pm 4.2$ | $761 \pm 201$ | 10.1 |
| R292Q<sup>†</sup> | $13.0 \pm 1.6$ | $14.6 \pm 4.6$ | $887 \pm 301$ | 11.8 |
| H296P | $14.7 \pm 0.9$ | $7.04 \pm 1.4$ | $2090 \pm 424$ | 27.7 |
| H296R<sup>†</sup> | $17.8 \pm 2.9$ | $30.8 \pm 9.9$ | $579 \pm 208$ | 7.67 |
| P300L | $14.0 \pm 0.7$ | $1.66 \pm 0.4$ | $8460 \pm 1880$ | 112 |
| T327R | $15.9 \pm 0.8$ | $1.69 \pm 0.4$ | $9460 \pm 2300$ | 125 |
| I361T | $17.4 \pm 0.7$ | $2.02 \pm 0.4$ | $8580 \pm 1600$ | 114 |
| E368G | $17.6 \pm 1.0$ | $1.96 \pm 0.5$ | $8990 \pm 2340$ | 119 |

<sup>1</sup> Mutant CKB assays were performed with $n = 3$ , while WT CKB was performed $n = 8$ , and data were globally fit to the Michaelis-Menten equation. Fitted kinetic parameters are shown with standard error. The data obtained from R132H better fit a linear regression, thus $k_{cat}$ and $K_m$ values could not be determined and are reported as N/A (not applicable). <sup>†</sup>: mutants that displayed reduced formation of TFV-DP in the in vitro activity assay.

R172P and R172G had similar Km values to WT, yet compared to each other showed a slight deviation in their kcat values, revealing a mutation-specific change in enzymatic turnover. H296R displayed a Km 15 times larger than WT and 4.4 times larger than H296P, indicating the presence of mutation-specific alterations in substrate binding affinity. Of particular interest, the R132H mutation better fit a linear regression, thus only the kcat/Km parameter could be determined, which showed to be only 2.10% of WT. Additionally, D268H resulted in a catalytic efficiency over 2 times larger than WT, driven by a Km half that of WT, yet displayed no significant change in ATP or TFV-DP formation. Interestingly, the C74S mutation had a catalytic efficiency 1.66 times larger than WT, also as a result of a lower Km, but exhibited no significant change in ATP formation and a 79.9% decrease in TFV-DP formation. These data are curious, but as mentioned previously this particular mutation demonstrates substrate-dependent activity (**Figure 2**), thus these apparent contradicting findings are a function of the fact that more than one substrate was used in our analyses. The enzyme kinetics assay employed here exploited the reverse canonical reaction, where ATP is binding and being dephosphorylated, therefore, data obtained using this assay are limited to examining the kinetics of this reaction. Of note, CKB exists as a dimer and WT-mutant heterodimers can display a dominant negative phenotype.^39^ The in vitro experiments reported here model a homozygous genotype, therefore further studies are warranted to characterize heterozygosity.

Lastly, differential scanning fluorimetry was employed to examine the impact of these mutations on the thermal stability of the enzyme. Five naturally occurring mutations (C74S, R96P, R132H, R236Q, and H296P), as well as C283S, displayed double peaks in their melting curves. The melting temperatures of the first peaks in C74S, R96P, and H296P had significantly lower melting temperatures than that of WT. Further, R172P showed prominent shouldering prior to the dominant peak, which exhibited a significantly lower melting temperature than WT **(Figure 3)**. These unique melting curves imply mutation-induced dimer or local domain instabilities. Arg172 falls near the dimer interface, therefore we predict the shouldering is a result of dimer melting. Additionally, a double peak was observed for all three mutations located in the phosphocreatine binding pocket (C74S, R96P, and C283S), suggesting mutations in this region may lead to local domain instability. These data reveal regions that may be sensitive to body temperature fluctuations caused by mutation-induced disorder.

**Figure 3.**
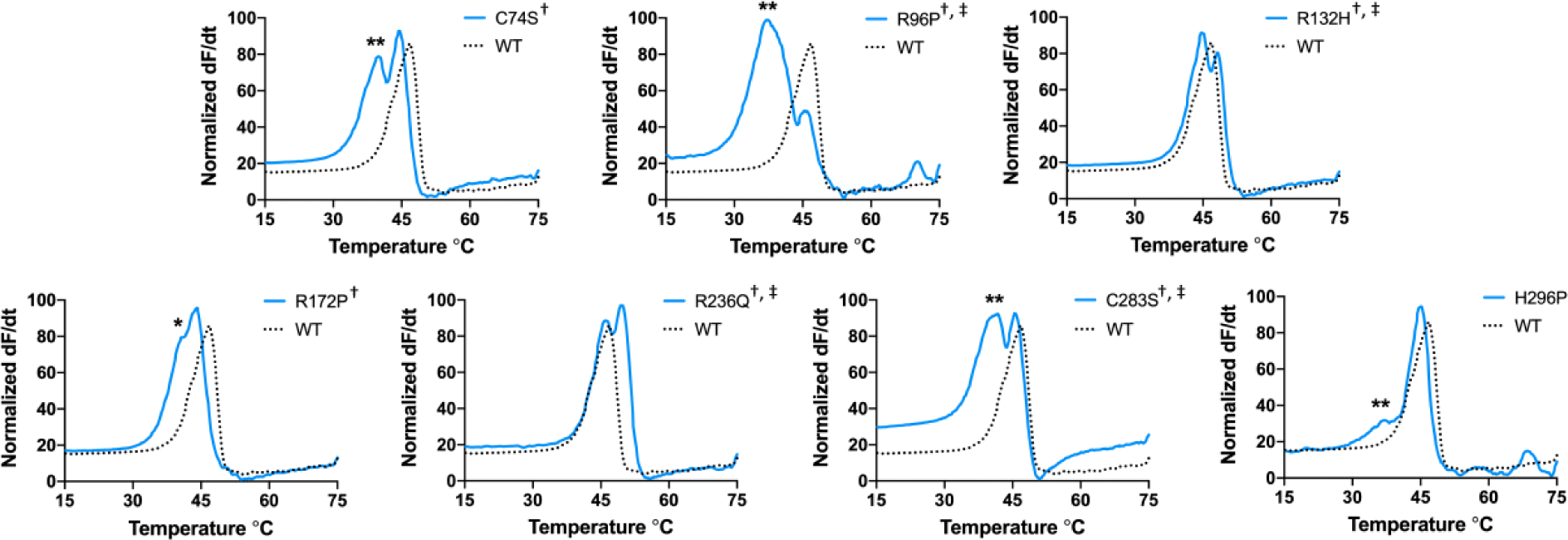
Differential scanning fluorimetry utilizing SYPRO orange dye reveals double peaks or shouldering of six naturally occurring mutants and the C283S negative control mutant (blue, solid) when compared to WT (black, dotted). Statistical analyses of melting temperatures (peak maxima) were performed using a one-way Brown-Forsythe and Welch ANOVA with Dunnett’s multiple comparisons test; P-values < 0.05*, 0.01**. ^†^: mutants that displayed reduced formation of TFV-DP in the in vitro activity assay, ^‡^: mutants with < 20% the ATP dephosphorylation catalytic activity of WT.

### CRISPR/Cas9-mediated Generation of *Ckb^tm1Nnb^* Mice

To examine the contribution of CKB in the formation of TFV-DP in vivo, CRISPR/Cas9 was used to target the start codon of the *Ckb* gene in C57BL/6J mice. The *Ckb* gene from founder mice was PCR amplified and Sanger sequenced, and mice with predicted deleterious mutations were outbred to WT. The most productive male breeder possessed two mutations on the same allele, a 55 base pair deletion that spanned 12 base pairs of intron 1 and 43 base pairs of exon 2, including the start codon, and a 28 base pair deletion beginning 134 base pairs into exon 2 **(Figure 4 A,B)**. mRNA extracted from the brains of WT and homozygous mice (n= 3) was reverse transcribed into cDNA and PCR amplified. Sanger sequencing revealed that the two base pairs (AG) following the 55 base pair deletion acted as a new 3’ splice site acceptor. If this mRNA transcript is translated into protein, translation would begin 22 base pairs into the mutant exon 2 and would code for just five amino acids **(Figure 4C)**. This allele was used to generate the homozygous *Ckb^tm1Nnb^* strain. Subsequent genotyping was completed by PCR amplification of the mutated region of the gene (**Figure 4D**). Next, quantitative PCR determined that *Ckb* expression is significantly lower in the brains of *Ckb^tm1Nnb^* mice and total protein knockout was confirmed in brain lysates by immunoblot **(Figure 4E,F)**.

**Figure 4.**
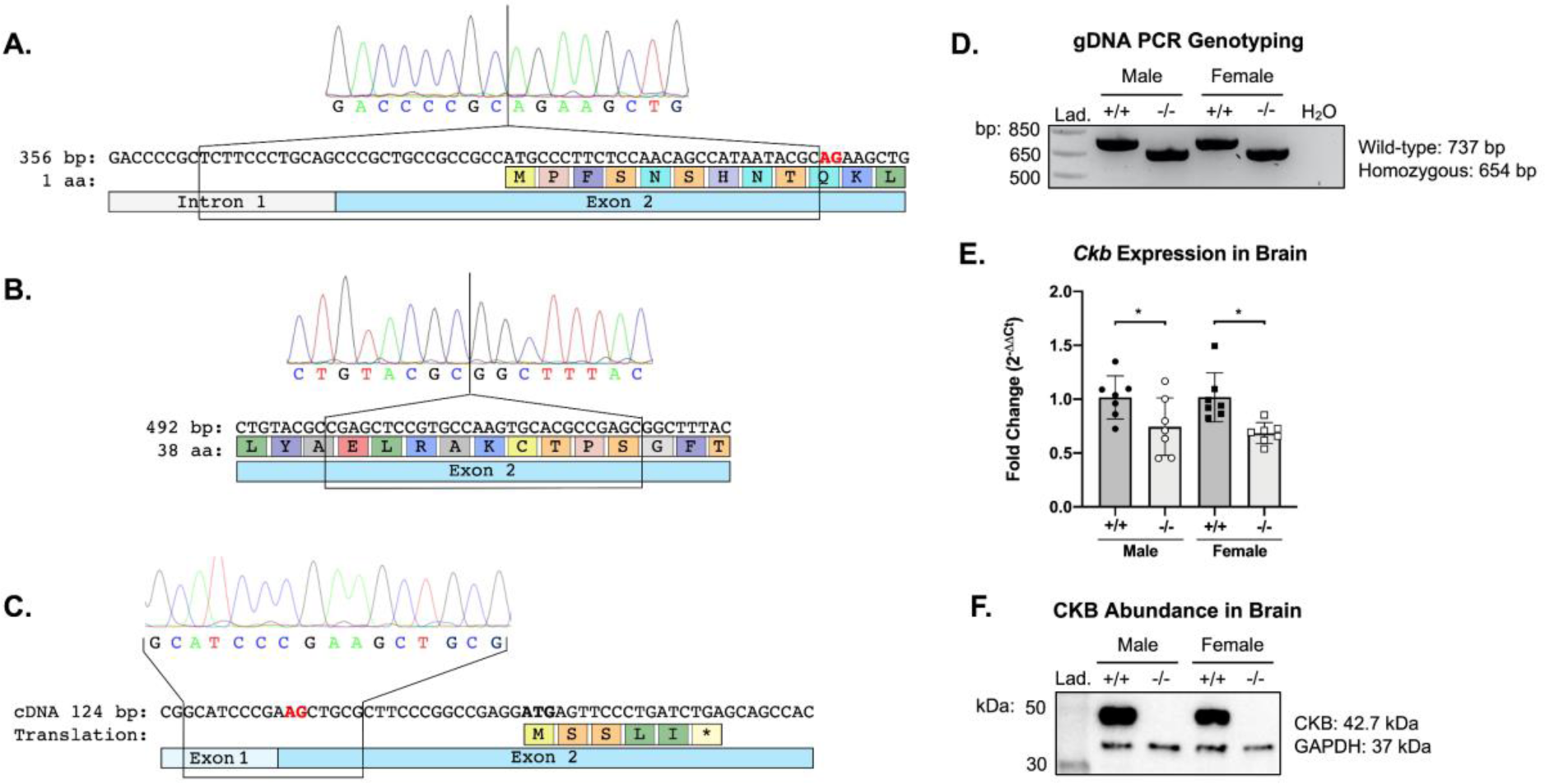
*Ckb* knockout mice were generated using CRISPR/Cas9. The most productive male breeder possessed two mutations on the same allele, (**A**) a 55 bp (base pair) (**B**) a 28 bp deletion, located at the guide RNA oligonucleotides target sites. mRNA extracted from brains of homozygous mice was reverse transcribed into cDNA and the first 5 exons were amplified by PCR. (**C**) Sanger sequencing revealed a new 3’ splice site, a new out-of-frame start codon, and the absence of any mutation-induced alternative splicing. If this transcript is translated, a protein of just 5 amino acids would be produced. (**D**) Genomic DNA (gDNA) was amplified by PCR and run on an agarose gel to genotype subsequent litter. (**E**) Using quantitative PCR, *Ckb* expression was determined to be significantly lower in homozygote mice (n = 7). (**F**) Immunoblotting for CKB shows a total loss of CKB protein in homozygote brain lysates. Statistical analyses of *Ckb* expression were performed on the Log(Fold Change) using an ordinary one-way ANOVA with Sidak’s multiple comparisons test; P-values < 0.05*. Lad. = ladder.

### TFV-DP Formation is Reduced in Brain Lysates

Active protein lysates from the brains of WT and *Ckb^tm1Nnb^* mice were obtained by gently homogenizing full brains in assay buffer and briefly centrifugating to remove cellular debris. Lysates were incubated in the presence of TFV-MP and a mix of ATP, phosphocreatine, and phosphoenolpyruvate. Resulting TFV metabolites were extracted and detected using uHPLC-MS. *Ckb^tm1Nnb^* male lysates formed 70.5% less TFV-DP than WT male lysates (n = 3, p<0.0001) and *Ckb^tm1Nnb^*female lysates formed 77.4% less TFV-DP than WT female lysates (n = 4 and 3, respectively; p<0.001) (**Figure 5A**). From this, we predict that CKB is the main kinase contributing to the formation of TFV-DP in brain. However, TFV-DP is still being formed in the *Ckb^tm1Nnb^*lysates, implicating the presence of other enzymes capable of contributing to this reaction. Additionally, WT female lysates formed 45.9% less TFV-DP than WT male lysates (n = 3, p<0.001) indicating the presence of sexual dimorphisms (**Figure 5A**). Because this assay measures TFV-DP formation in tissue lysates, we are unable to make conclusions regarding cell type- or region-specific differences or changes in blood brain barrier permeability between sexes or genotypes. However, it is known that TFV is phosphorylated in the human brain and reaches concentrations higher than other commonly prescribed nucleot(s)ide reverse transcriptase inhibitors.^41^

**Figure 5.**
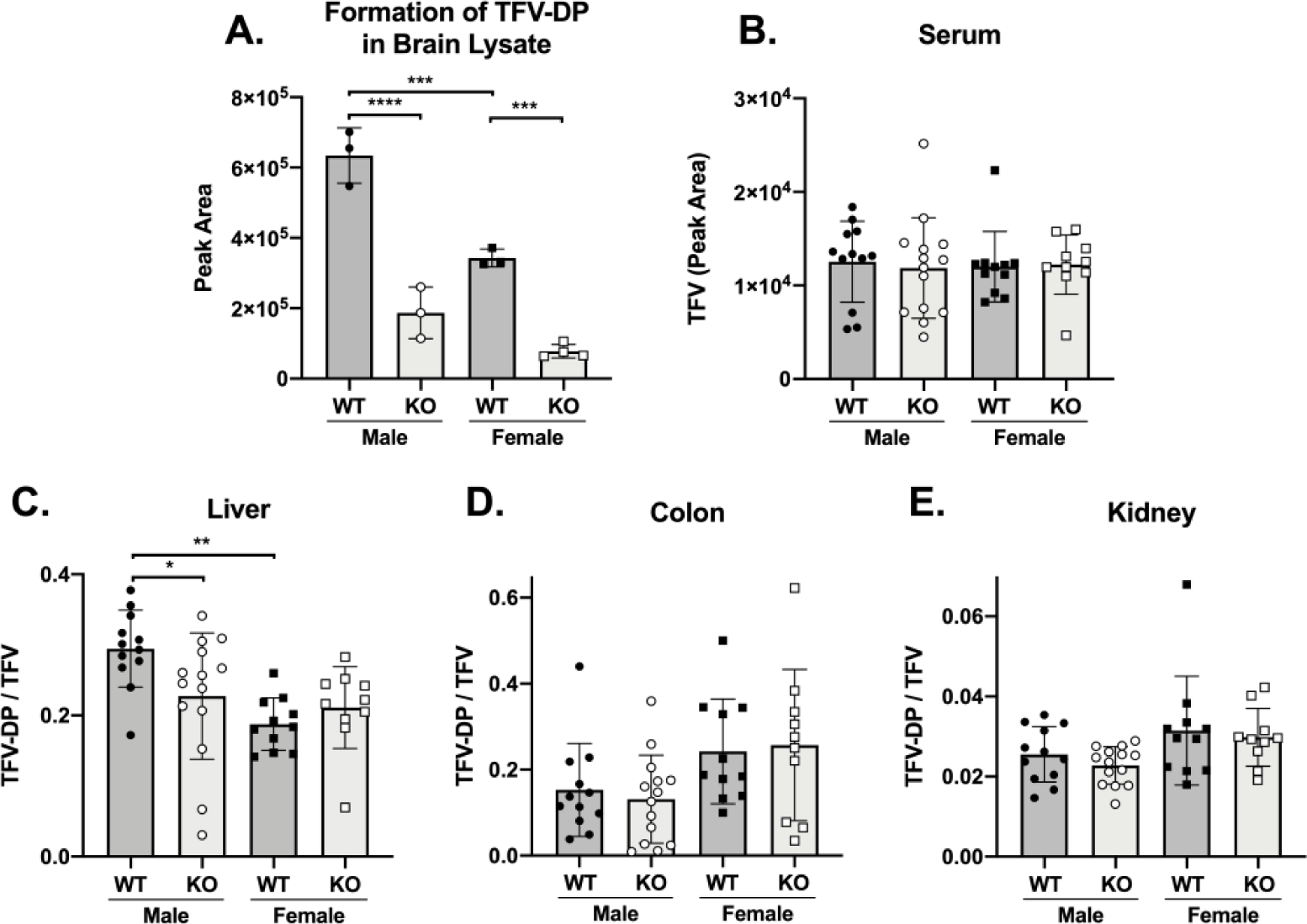
Tissue-specific TFV activation in wild-type (WT) and *Ckb^tm1Nnb^* (KO) mice. (**A**) Full brain lysates were incubated in the presence of TFV-MP, ATP, phosphocreatine, and phosphoenolpyruvate (n = 3 or 4). Formation of TFV-DP was detected using uHPLC-MS. Wild-type and KO mice were dosed orally via drinking water and TFV metabolites were extracted from harvested tissues and detected by uHPLC-MS. (**B**) TFV is the only metabolite present in serum and shows no change between sexes or genotypes. TFV activation (TFV-DP/TFV) was calculated for liver, colon, and kidney samples. (**C**) In liver, KO males and WT females have significantly lower TFV activation than WT males. (**D,E**) No change in TFV activation was observed in colon or kidney. Statistical analyses were performed using an ordinary one-way ANOVA with Sidak’s multiple comparisons test; P-values < 0.05*, 0.01**, 0.001***, 0.0001****.

### TFV Activation is Reduced in *Ckb^tm1Nnb^* Liver but Not in Colon or Kidney

Next, to understand where and to what extent CKB contributes to TFV activation in vivo, mice were dosed orally (60mg/kg/day) with tenofovir disoproxil fumarate/emtricitabine for 14 days. TFV metabolites were extracted from serum, liver, colon, and kidney tissue and detected by uHPLC-MS. The only metabolite found in serum was TFV, which showed no difference between dosed groups, suggesting TFV bioavailability is unchanged in *Ckb^tm1Nnb^* mice (**Figure 5B**). To assess TFV activation in liver, colon, and kidney, we calculated the ratio of the area under the TFV-DP peak curve to the area under the TFV peak curve for each sample. In liver, TFV activation was decreased 22.8% in *Ckb^tm1Nnb^* male mice compared to WT male mice (n=14 and 12, respectively; p<0.05) (**Figure 5C**). No difference was observed between WT and *Ckb^tm1Nnb^* female liver samples. However, WT females had 36.3% less TFV activation than WT male livers signifying the presence of sexual dimorphisms. No differences in TFV activation were observed in colon or kidney (**Figure 5 D,E**). In colon, it has been previously demonstrated that another creatine kinase, CKM, is the primary enzyme that catalyzes the formation of TFV-DP, thus no change in TFV activation was expected.^17^ A similar redundancy of other TFV activating enzymes may also explain the lack of change in kidney across genotypes. No significant differences in TFV activation were present in liver, colon, or kidney of WT and *Ckb^tm1Nnb^*mice dosed for 24 hours (data not shown).

### Enzymes Involved in TFV-DP Formation are Differentially Abundant in *Ckb^tm1Nnb^* Brains

Proteomic analyses were conducted on brain samples of male and female WT and *Ckb^tm1Nnb^* mice to better understand the tissue-specific effect of knocking out CKB, with a particular interest in nucleotide and small molecule kinases. Full brains were homogenized and proteins were prepared for mass spectrometry-based data independent acquisition proteomic analysis. Using the directDIA default workflow adjusted to identify only proteotypic peptides, Spectronaut identified 5,818 proteins across brain samples, from which we curated a list of 123 nucleotide and small molecule kinases (**Supplementary Table 3**). The MS2 area for each peptide of a protein was summed to provide a protein quantity. Protein quantities were uploaded into the SimpliFi interpretation program where statistical analyses were performed (public access to view the data in SimpliFi is available at https://tinyurl.com/CKBbrain). Of the 123 nucleotide and small molecule kinases, 57 proteins have a q-value<0.05 in at least one comparison, chemically modify or use ATP as a substrate, and have a reaction compatible with the structure of TFV metabolites (**Supplemental Table 4**). For the remainder of this work, proteins detected by proteomics will be referred using their unique gene ID.

Interestingly, enzymes that show activity toward TFV-DP formation displayed differential abundances in brain. Ckm, Nme1, and Pkm, which can phosphorylate TFV-MP to TFV-DP, showed reduced abundance in *Ckb^tm1Nnb^*mice as compared to WT (**Figure 6A**).^17,42^ The abundance of Ckb in WT female brains was 60% that of WT males, and the abundance of Ckm in *Ckb^tm1Nnb^* brains was less than 10% that of WT brains of the same sex. *Ckb^tm1Nnb^* brains had a significantly lower abundance of Nme1 and Pkm when compared to WT brains of the same sex. Further, Entpd1, which is able to dephosphorylate TFV-DP to TFV, was increased in WT female and *Ckb^tm1Nnb^*male brains compared to WT male brains.^43^ Together, the differential abundances of these enzymes likely compound the loss of Ckb, resulting in the significant reduction in TFV-DP formation we observed in *Ckb^tm1Nnb^*brain lysates.

**Figure 6.**
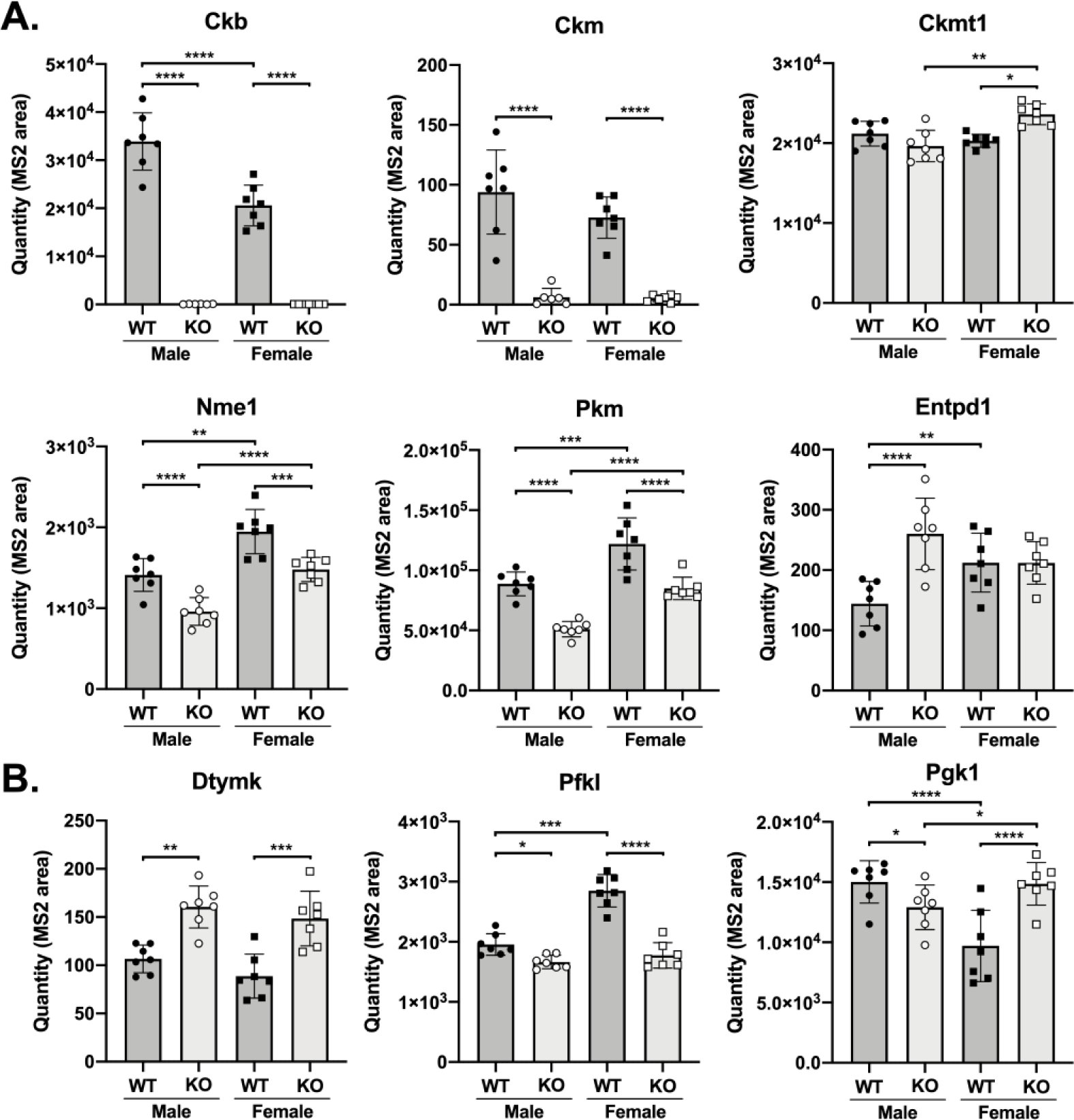
Proteomic analyses of brain lysates reveal significant changes in the abundance of nucleotide and small molecule kinases. (**A**) Creatine kinases (Ckb, Ckm, and Ckmt1), Nme1, and Pkm are able to phosphorylate TFV-MP to TFV-DP, while Entpd1 is able to dephosphorylate TFV-DP to TFV. (**B**) Dtymk is a nucleotide kinase that reversibly transfers a phosphate to ATP, Pfkl and Pgk1 transfer a phosphate group from ATP to small molecules. Statistical analysis is completed using SimpliFi. P-values<0.05*,<0.01**,<0.001***,<0.0001****.

To assess the abundance changes in nucleotide and small molecule kinases beyond those with known TFV activity, the 57 proteins listed in **Supplemental Table 4** were surveyed further. We found that Dtymk and Pfkl displayed differential abundances that align with the TFV-DP formation data from the brain lysate in vitro assay (**Figure 6B**). The nucleotide kinase Dtymk, which transfers a phosphate from ATP to dTMP, was 1.5 and 1.7 times higher in *Ckb^tm1Nnb^* male and female mice compared to WT mice of the same sex, respectively.^44^ If Dtymk can bind TFV-DP, dephosphorylation may occur and reduce TFV-DP levels, but additional studies are required to determine whether TFV-DP is a Dtymk substrate. Pfkl, the rate limiting enzyme in glycolysis that transfers a phosphate from ATP to fructose-6-phosphate to form fructose-1,6-bisphosphate, was most abundant in WT female brains, approximately 1.5 times greater than WT male brains and 1.6 times greater than *Ckb^tm1Nnb^*female brains. However, no sex-dependent change was observed between male and female *Ckb^tm1Nnb^* brains. Pgk1, a small molecule kinase that transfers a phosphate from ATP to phosphoglycerate, is of particular interest because its canonical reaction is inhibited by TFV-MP, yet displays no activity toward TFV-MP phosphorylation.^45^ Further, Pgk1 is known to contribute to the activation of emtricitabine, another nucleotide analog used to treat/prevent HIV infection.^20^ The abundance of Pgk1 in WT female brains was significantly lower than in WT male brains, while *Ckb^tm1Nnb^*female brains had a significantly higher abundance of Pgk1 than *Ckb^tm1Nnb^*male brains. Together, these data indicate that loss of CKB in the brain alters the abundance of nucleotide and small molecule kinases, which may affect physiological processes and the activation of other nucleotide analogs.

### Creatine Kinases are Not Detected in Liver but Nucleotide and Small Molecule Kinases are Differentially Abundant

To examine the differential abundance of proteins between male and female WT and *Ckb^tm1Nnb^* livers, proteins were identified using mass spectrometry-based data independent acquisition proteomics (public access to view the data in SimpliFi is available at https://tinyurl.com/CKBliver). We identified 4,720 proteins across liver samples, from which we curated a list of 100 nucleotide and small molecule kinases (**Supplemental Table 3**). Of the 100 nucleotide and small molecule kinases, 61 proteins have a q-value<0.05 in at least one comparison, chemically modify or use ATP as a substrate, and have a reaction compatible with the structure of TFV metabolites (**Supplemental Table 5**). Surprisingly, no creatine kinase enzymes were identified using our proteomic analyses, even though mRNA of *CKB*, *CKMT1*, and *CKMT2* have been detected in human liver biopsies.^21^ Thus, we postulate that the reduction in TFV activation in *Ckb^tm1Nnb^*male livers may be due to an indirect effect of the CKB knockout. We propose that the effects of the full body knockout can propagate across tissues, resulting in differentially abundant proteins even in tissues where CKB is not present. For example, we observe that adenylate kinase and pyruvate kinase isoenzymes are differentially abundant (**Figure 7A**). While the activity of Ak3 and Ak4 toward TFV is unknown, due to high homology to Ak2, we hypothesize they may also contribute to the phosphorylation of TFV to TFV-MP.^46^ In *Ckb^tm1Nnb^*livers, Ak3 and Ak4 were significantly more abundant than WT livers of the same sex. In contrast, Pklr and Pkm were significantly lower in abundance in *Ckb^tm1Nnb^*male livers, compared to WT male livers. Pklr was 1.48-fold greater in WT males than WT females, yet WT females had 1.76-fold more Pkm than WT males. These abundance differences may explain the sexual dimorphism in TFV activation between WT mice if TFV-MP has an equal or higher affinity for Pklr than Pkm. Multiple nucleotide kinases and small molecule kinases with unknown activity toward TFV displayed a fold change > 1.5 in at least one comparison. Interrogating these data for sexual dimorphisms, Adk and Cmpk1, which reversibly transfer a phosphate to ATP, were significantly lower in WT female livers. Nucleotide phosphatases Enpp3 and Entpd5 were significantly increased in *Ckb^tm1Nnb^* male livers, which may negatively regulate TFV activation (**Figure 7B**). Further, Idnk, Mvk, Pgk1, and Tkfc transfer a phosphate group from ATP to small molecules and are differentially abundant across genotypes and sexes, following a similar trend to the sexual dimorphisms in TFV activation. If TFV metabolites are substrates of these enzymes, they may also contribute to the regulation of TFV activation.

**Figure 7.**
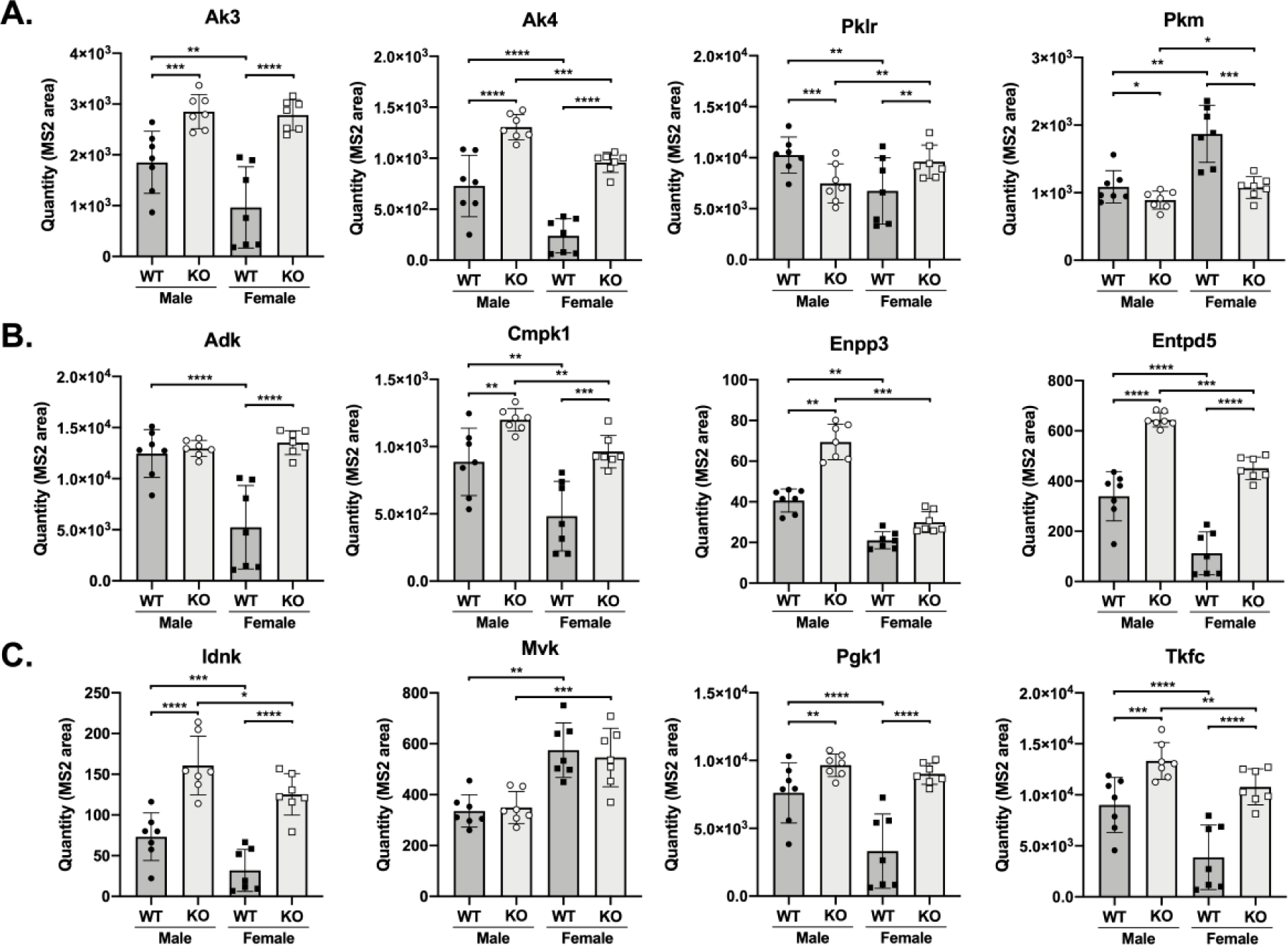
Proteomic analyses of liver lysates reveal significant changes in the abundance of nucleotide and small molecule kinases. (**A**)Adenylate kinases (Ak3 and Ak4) and pyruvate kinases (Pklr and Pkm) are able to phosphorylate TFV to TFV-MP and TFV-MP to TFV-DP, respectively. (**B**) Adk and Cmpk1 are nucleotide kinases that reversibly transfer a phosphate to/from ATP, while Enpp3 and Entpd5 are nucleotide phosphatases. (**C**) Idnk, Mvk, Pgk1, and Tkfc transfer a phosphate group from ATP to small molecules. Statistical analysis is completed using SimpliFi. P-values < 0.05*, < 0.01**, < 0.001***, < 0.0001****.

## CONCLUSION

Together, our data suggest that naturally occurring mutations in CKB can diminish the formation of TFV-DP. We provide evidence that CKB catalyzes the formation of TFV-DP in brain and that TFV activation in the liver can be reduced if CKB function or abundance is decreased. Not only do we expect CKB to directly contribute to the activation of other nucleot(s)ide analogs, our proteomics data reveal that loss of CKB disrupts the proteomic landscape of other nucleotide and small molecule kinases. We predict this may alter the activation of certain co-administered nucleotide analogues, such as emtricitabine, that require phosphorylation by these enzymes. In conclusion, we hypothesize that individuals possessing missense mutations or diseases that limit the catalytic activity of CKB may have lower levels of TFV-DP in the brain and liver, thereby potentially impacting the pharmacology of TFV in the treatment of HIV and HBV infections.

## MATERIALS AND METHODS

### Selection of Naturally Occurring Mutations

Ensembl’s Variant Effect Predictor was used to identified 258 genetic variants in dbSNP build 151.^47^ Missense mutations predicted to be damaging by SIFT and PolyPhen scoring that were located in the phosphoryl transfer domain, phosphocreatine or ADP binding pockets, or at the dimer interface and had an allele frequency of approximately 0.00001 were selected for further analysis.^40,48,49^ Several missense mutations found on the external surface of the protein were included as controls. CKB C283S mutation was selected as a negative control, as it is functionally null.^39^

### Vector Cloning and Site-Directed Mutagenesis

Human CKM, CKB, and creatine kinase U-type (CKMT1) cDNA (Origene, Rockville, MD) were cloned into pET-30a vectors (Sigma-Aldrich, St. Louis, MO) as previously described.^19^ Site-directed mutagenesis was performed using the QuikChange Lightning Site-Directed Mutagenesis Kit (Agilent, Santa Clara, CA) according to the manufacturer’s protocol. Mutagenesis primers were designed using the QuikChange Primer Design web-based tool (Agilent) and purchased from Integrated DNA Technologies (**Supplemental Table 1**). The sequence of each mutant CKB vector was confirmed by Sanger sequencing using the Johns Hopkins University Synthesis and Sequencing Facility.

### Recombinant Protein Expression and Purification

Vectors were transformed into BL21(DE3) competent *Escherichia coli* (Agilent) and a starter culture was grown at 37°C overnight in 10 ml Luria Broth (LB) containing 50 μg/ml kanamycin. The starter culture was added to 1 liter of autoclaved LB containing 50 μg/ml kanamycin and incubated at 37°C for 3 hours while shaking at 220rpm. Once the OD600 reached 0.6-0.8, isopropyl β-D-1-thiogalactopyranoside was added to achieve a final concentration of 1mM to induce expression. After 3 hours, the culture was pelleted by centrifugation at 4,000 x *g* for 7 minutes and stored at - 80°C. Protein purification was performed as described previously.^19^ In summary, bacterial pellets were lysed, and supernatants were incubated with cobalt resin (Takara) for 1 hour at 4°C. The column underwent a wash of low- and high-salt buffers and His-tagged CKB enzymes were eluted with increasing concentrations of imidazole. Absorbance at 280 nm was determined using a Take3 microvolume plate in a plate reader (BioTek Synergy HT) and protein concentration was calculated using a path length of 0.05 cm and an extinction coefficient of 35,275 M^-1^ cm^-1^. On average, 10.9 mg of protein was purified per purification. Protein purity was determined through SDS-PAGE with Coomassie blue stain.

### In Vitro Formation of TFV-DP

Assays were conducted as previously reported, with the adjustment that reactions were quenched with 200μl of ice-cold 100% methanol (n=3).^19^ Formation of TFV-DP was detected by mass spectrometry as previously described using a Dionex Ultimate 3000 uHPLC (Thermo Fisher) coupled to a TSQ Vantage triple quadrupole mass spectrometer (Thermo Fisher).^19^ Column temperature was maintained at 40°C. Peak area of TFV-DP formed by mutant enzymes was normalized to the average WT peak area of TFV-DP. Statistical analyses were performed by GraphPad Prism (version 8.4.3) using a one-way Brown-Forsythe and Welch ANOVA with a Dunnett’s multiple comparisons test comparing each mutant CKB to wild-type CKB.

### In Vitro Formation of ATP

Assays were performed and the formation of ATP was detected as previously described, with the exception that 200μl of ice-cold 100% methanol was used to quench the reaction.^19^ Statistical analyses were performed as mentioned above.

### ATP Dephosphorylation Kinetics

An enzyme coupled system was established to measure enzyme kinetics. The reaction utilizes the reverse canonical reaction of CKB producing ADP, which is then used in the pyruvate kinase/lactate dehydrogenase system resulting in the conversion of NADH to NAD^+^. Assays and analyses were performed as previously described, with minor adjustments to ATP concentration series.^19^ Briefly, purified protein was prewarmed at 37°C for 5 minutes in assay buffer (75 mM HEPES pH 7.5, 5 mM MgCl2, 50 mM KCl, 2 mM DTT, 1 mM phosphoenolpyruvate, 20 mM creatine, 20 nM CKB, 66 units/ml pyruvate kinase (rabbit), 105 units/ml lactate dehydrogenase (rabbit), 400 nM NADH), after which the reaction was initiated by addition of ATP (final concentrations: 50 mM, 35 mM, 20 mM, 10 mM, 5 mM, 2.5 mM, 1.25 mM, 0.625 mM, 0.156 mM, and 0 mM). Reaction progression was monitored by absorbance at 340 nm (loss of NADH) every 10 seconds for 3 min. Initial rates were calculated using a linear fit (Microsoft Excel) and plotted against the concentration of ATP. Data was globally fit to the Michaelis-Menten equation and kinetic parameters were obtained using GraphPad Prism (version 8.4.3). The R132H CKB mutant better fit a linear regression than the Michaelis-Menten equation, thus only the kcat/Km kinetic parameter could be determined.

### Differential Scanning Fluorimetry

Assays were performed as previously described.^19^ Briefly, purified protein and SYPRO Orange dye (final concentrations: 5 μM and 25X, respectively) were gently mixed in assay buffer (75 mM HEPES pH 7.5, 5 mM MgCl2, 50 mM KCl, 2 mM DTT). Using an Eppendorf Mastercycler RealPlex4 instrument, fluorescence was detected at 580 nm every 0.2-0.3°C as temperature increased from 15°C to 75°C at a ramp speed of 2°C/min. The first derivative of the raw fluorescence data was calculated by RealPlex software (Eppendorf). Three technical replicates of each protein were run in parallel and averaged to obtain a melting curve. Each curve was normalized to its highest value. This was completed four times, and the melting temperatures were determined as the maxima in each replicate. After all replicates were complete, melting curves of the same mutant were averaged and qualitatively compared to WT. Statistical analysis comparing each melting temperature to the melting temperature of WT were performed as described above.

### CRISPR-Cas9 Targeting Strategy

Two gRNA were designed to target upstream and downstream of the *Ckb* start codon in C57BL/6J mice (gRNA1: GCGGGCUGCAGGGAAGAGCGUUUUAGAGCUAUGCU and gRNA2: CGUGCCAAGUGCACGCCGAGGUUUUAGAGCUAUGCU). RNA was purchased from Integrated DNA Technologies. Preparation, pronuclear injections, and surgeries were completed by the Johns Hopkins Transgenic Core Laboratory.

### Genotyping and Sequencing of *Ckb*

Genomic DNA was extracted from tail snips using the DNeasy Blood and Tissue Kit (Qiagen) following the manufacturer’s protocol for purifying total DNA from animal tissues. DNA was stored at 4°C. PCR reactions contained 1X Phusion High-Fidelity PCR Master Mix (New England BioLabs Inc.), 0.5 μM primers (forward: CATCCATCGCCCCTGCTTCGTC, reverse: CGGGCGACGAGGAGAGTTACGA), and 100 ng DNA in a 50 μl volume. Reactions were completed using an Eppendorf Mastercycler pro with vapo.protect lid with the following cycling conditions: denaturation at 98°C for 30 sec, 35 cycles of 98°C (10 sec), 73°C (30 sec), and 72°C (30 sec), and a final extension at 72°C for 5 min. Reactions were mixed with Gel Loading Dye Blue (New England BioLabs Inc.) and ran on a 2% agarose gel with 1X SYBR Safe DNA gel stain (Invitrogen) for 45 min at 100V. Gels were imaged under UV light (BioRad). Samples with apparent shifts in PCR band molecular weight were prepared for Sanger sequencing. Samples were loaded into Ultracel-10 regenerated cellulose membrane centrifuge filters (Millipore Sigma) with 400 μl H2O and centrifuged at 14,000 x *g* for 10 min. This was repeated twice. The filter was inverted into a clean tube and centrifuged at 17,000 x *g* for 3 minutes. The resulting PCR product was diluted to 10 ng/μl and mixed with 4 μM nested primers (forward: GGGAACTTGGGATGCGCTGGAC, reverse: CTTCCCTGAACCTTCGGTGGGC). Sanger sequencing was completed by the Johns Hopkins University Synthesis and Sequencing Facility.

### Quantitative PCR

To determine the expression of the *Ckb* gene across sexes and genotypes, mRNA was extracted from WT and *Ckb^tm1Nnb^* mice (n=7). Harvested brains were placed in 700μl TRIzol solution (Ambion) and lysed by bead milling (Beadbug Benchmark Scientific). Lysates were then split into two tubes and frozen at -80°C. Additional TRIzol was added to each lysate to reach 1000 μl prior to the addition of 200 μl chloroform. Samples were centrifuged at 12,000 x *g* at 4°C for 15 min and the aqueous layer was removed. Next, 200 μl 100% ethanol was added dropwise while being gently vortexed and placed into a RNeasy Mini column (Qiagen), The remainder of the purification followed the manufacturer’s protocol. RNA was quantified using a Take3 microvolume plate (BioTek Synergy HT). Using an Eppendorf Mastercycler pro with vapo.protect lid, 2 μg of RNA was reverse transcribed using the High-Capacity cDNA Reverse Transcription kit (Applied Biosystems) following the manufacturer’s protocol. cDNA was diluted 1:2 in nuclease-free water and mixed with 500 nM forward and reverse primers and PowerUp SYBR Green Master Mix (Applied Biosystems). qPCR was conducted using an Eppendorf Mastercycler RealPlex4 instrument with the following cycling conditions: 50°C (2 min), 95°C (2 min), 40 cycles of 95°C (15 sec) and 60°C (1 min), then a hold at 4°C. Ct values were determined at fluorescence threshold of 1200. *Ckb* was normalized to *Gapdh* for relative quantification using the delta-delta Ct method. (*Ckb* forward: GGCGACGAGGAGAGTTACGACG; *Ckb* reverse: CATCGCCACCCTGCAGGTTGTC; *Gapdh* forward: CTCACTGGCATGGCCTTCCGTG; *Gapdh* reverse: CTTGGCAGGTTTCTCCAGGCG). Statistical analyses were performed on the Log(Fold Change) using an ordinary one-way ANOVA with Sidak’s multiple comparisons test.

### Animal Husbandry

All studies using mice were approved by Johns Hopkins Animal Care and Use Committee and have been carried out in accordance with the Guide for the Care and Use of Laboratory Animals as adopted and promulgated by the U.S. National Institutes of Health. Animal housing rooms remained at 72°C with 42% humidity. Lights turned on at 6:30 am and off at 9:00 pm. Animals had unrestricted access to water and Teklad Global 18% Protein Extruded Rodent Diet chow (Envigo).

### Immunoblot of CKB

To verify protein knockout, immunoblot analyses were performed using brain samples from WT and *Ckb^tm1Nnb^*sibling littermates. Animals were euthanized via isoflurane overdose, followed by cervical dislocation. Brains were harvested, frozen, and stored at -80°C until analysis. Each sample was homogenized using a bead mill (BeadBug Benchmark Scientific) in 500 μl cell lysis buffer: 1X cell lysis buffer (Cell Signaling Technology), 500 μM phenylmethylsulfonyl fluoride (Thermo Fisher), and 1X Halt Protease Inhibitor Cocktail (Thermo Scientific). Cellular debris was pelleted by centrifuging at 13,000 x *g* for 20 min at 4°C. Total protein was quantified by absorbance at wavelength 280 nm using a Take3 microvolume plate (BioTek Synergy HT) and 1 μg was boiled for 10 min in 1x Laemmli Sample Buffer (BioRad) with 5% 2-mercaptoethanol. Samples were then run on a 4-15% polyacrylamide gel (BioRad) at 165 V for 40 min. Proteins were transferred to a nitrocellulose membrane using the iBlot2 transfer device (Life Technologies Inc.) and the membrane was cut to allow blotting for both CKB and GAPDH. The membrane was blocked using 5% non-fat dry milk (BioRad) in TBST (Sigma-Aldrich) for 1.5 hour at room temperature. The membrane was incubated with 1:10,000 anti-CKB antibody (ab126418, Abcam) and 1:1,000 anti-GAPDH (#5174, Cell Signaling Technologies) in 5% bovine serum albumin in TBST at 4°C overnight. The membrane was shaken three times for 5 min in TBST then incubated with 1:3,000 anti-rabbit HRP-linked antibody (Cell Signaling Technologies) for 1 hour at room temperature. After incubation, the membrane was shaken in TBST for 5 min, three times then incubated for 2.5 min in West Dura solution (Thermo Scientific). Blots were imaged using a chemiluminescence imager (BioRad).

### TFV-DP Formation In Vitro

To examine the extent TFV-DP formation is altered in *Ckb^tm1Nnb^*mice, full brains were harvested from 12 week old WT (3 males, 3 females) and *Ckb^tm1Nnb^* (3 males, 4 females) mice. Brains were weighed and placed into bead mill tubes containing 1 ml of assay buffer (75 mM HEPES pH 7.5, 5 mM MgCl2, 50 mM KCl, and 2 mM DTT) for every 200 mg of tissue. Brains were homogenized using a bead mill (Beadbug Benchmark Scientific) for 30 sec at 4,000 rpm and resulting lysates were centrifuge at 3,000 x *g* at 4°C for 10 min. Total protein was quantified by absorbance at wavelength 280 nm using a Take3 microvolume plate (BioTek Synergy HT). In assay buffer, 50 μg of total protein and 100 μM TFV-MP were incubated for 5 min at 37°C, after which reactions were initiated with the addition of 500 μM ATP, 500 μM phosphocreatine, and 500 μM phosphoenolpyruvate. The reaction proceeded for 30 min, at which point 200 μl of ice cold methanol quenched the reaction. Samples were centrifuged at 10,000 rpm at 4°C for 10 min and supernatants were dried under vacuum centrifugation (Eppendorf) overnight. Samples were resuspended in 30 μl mobile phase A, centrifuged for 8 min at 13,500 x *g* for 8 min and the supernatant was injected into a uHPLC-MS as described above. Statistical analyses were performed by GraphPad Prism using an ordinary one-way ANOVA with a Sidak’s multiple comparison’s test comparing WT male/female, *Ckb^tm1Nnb^* male/female, WT male/*Ckb^tm1Nnb^* male, and WT female/*Ckb^tm1Nnb^*female.

### TFV-DP Formation In Vivo

To determine the in vivo contribution of CKB in catalyzing the formation of TFV-DP, WT (12 male, 11 female) and *Ckb^tm1Nnb^* (14 male, 10 female) mice were treated with tenofovir disoproxil fumarate (TDF) and emtricitabine (FTC), a drug combination used for PrEP. An interspecies allometric scaling factor of 12.3 was used to equate the human daily dose to that of mice, estimating a mouse daily dose of TDF of 60 mg/kg.^50^ The daily water intake of C57BL/6J mice is estimated to be 0.15 ml/g of body weight, thus crushed pill tablets were dissolved in drinking water at a TDF concentration of 0.400 mg/ml and an FTC concentration of 0.267 mg/ml. Due to the small scale of our colonies, mice were treated in three groups. In each group, 10 week ± 2 days old mice were treated for 14 days with unrestricted access to food or TDF/FTC water. After treatment, mice were euthanized by isoflurane overdose followed by cervical dislocation. Serum, liver, colon, and kidney were harvested, frozen, and stored at -80°C. Tissues were weighed and homogenized in 500 μl 70% methanol using a bead mill (Beadbug Benchmark Scientific). Resulting supernatants were dried down by vacuum centrifugation (Eppendorf) for 2 hr and resuspended in 1 μl of mobile phase A (H2O, 5 mM dimethylhexylamine) per 1 mg of tissue (kidney samples were resuspended in 1 μl mobile phase A per 2mg of tissue). To extract TFV from serum, 100 μl of serum was vortexed vigorously in 500 μl of 70% methanol and centrifuged at 13,000 x *g* at 4°C for 10 min. Supernatants were dried under vacuum centrifugation (Eppendorf) for 2 hr and resuspended in 40 μl mobile phase A. TFV-DP in each sample was detected as previously described using a Dionex Ultimate 3000 uHPLC (Thermo Fisher) coupled to a TSQ Vantage triple quadrupole mass spectrometer (Thermo Fisher).^19^ The only alteration was the addition of a Halo C18 90 Å, 2.7 μm, 2.1 x 5 mm guard column (Mac-Mod Analytical) to help preserve the column from this increased relative sample load, which was demonstrated to impart no noticeable change in assay detection limits (data not shown). TFV was detected in positive ion mode following the same instrument parameters as previously described and using a single-reaction monitoring scan with a mass to charge ratio transition of 288 to 176 with a collision energy of 35 V. The ratio of the peak area of TFV-DP to the peak area of TFV was used as a metric to define TFV activation in liver, colon, and kidney. Statistical analyses were performed as described in **TFV-DP Formation In Vitro**.

### Proteomic Analysis of Brain and Liver

To understand the change in the protein abundance of nucleotide and small molecule kinases, mass spectrometry-based data independent acquisition proteomics was performed. Proteins from brain and liver tissue of 12 week old WT and *Ckb^tm1Nnb^* male and female mice were extracted via homogenization in cell lysis buffer as described above (n=7 each group). Protein concentration was calculated using the Pierce BCA Protein Assay Kit (Thermo Scientific) following the manufacturer’s instructions. Next, proteins were digested into peptides using the S-Trap 96-well plate (ProtiFi) following the manufacturer’s instructions. In brief, 100 μg of protein from each sample were solubilized in 5% SDS, reduced with DTT (20mM), alkylated using iodoacetamide (40mM), acidified (1.2% phosphoric acid), trapped on column, then digested by 10 μg MS-grade trypsin (Thermo Scientific). Once peptides were eluted, they were dried under vacuum centrifugation (Eppendorf) overnight, resuspended in 100 μl 0.1% formic acid in H2O, and quantified using the Pierce Quantitative Colorimetric Peptide Assay kit (Thermo Scientific). Samples were diluted to 100 ng/μl and 2μl were injected by an EasyNLC 1200 (Thermo Scientific) nanoflow liquid chromatography system coupled to a timsTOF FleX mass spectrometer (Bruker). Mobile phase A was 0.1% formic acid in water and mobile phase B was 0.1% formic acid in 80% acetonitrile / 20% water. Peptides passed through an Acclaim PepMap C18 100Å, 3 μm, 75 μm x 2 cm trap column (Thermo Scientific) followed by separation on a PepSep C18 100Å, 1.5 μm, 75 μm x 25 cm (Bruker) at a flow rate of 200 nl/min using the following 1 hr gradient: 10% - 35% B from 0 - 47 min, 35% - 100% B from 47 – 55 min, 100% B from 55 min - 57 min, 100% - 5% B from 57 min - 58 min, and 5% B from 58 min - 60 min. Trap column equilibration used 9 μl at 3.0 μl/min flow rate, separation column equilibration used 12 μl at 3.0 μl/min flow rate. Additionally, 1 wash cycle of 20 μl and a flush volume of 100 μl were used. Peptides were ionized using the CaptiveSpray source, with a capillary voltage of 1500V, dry gas flow of 3 l/min and temperature 180°C. Data were acquired using a positive ion mode diaPASEF method with a mass range from 100 m/z to 1700 m/z and 1/K0 from 0.80 Vs/cm^2^ to 1.35 Vs/cm^2^ with 100 ms ramp time and 2 ms accumulation time. Full MS parameters have been published as “TIMS_40_shortgradient_DIA” and can be accessed at dx.doi.org/10.17504/protocols.io.x54v9p8zzg3e/v1. Resulting spectra were converted to .htrms file formats using HTRMS Converter (Biognosys) and uploaded into Spectronaut (Biognosys). Peptides were identified and quantified using the directDIA analysis default settings with the proteotypicity filter set to “Only Proteotypic”. Protein group quantifications were exported from Spectronaut and statistical analyses were performed by the internet-based SimpliFi platform (ProtiFi). All vendor original data and processed data has been made publicly available through the ProteomeXchange and MASSIVE public repositories.^51^ Data can be directly accessed through FTP at the following link: ftp://massive.ucsd.edu/MSV000091978/ or through ProteomeXchange.org as project PXD042296.

## ASSOCIATED CONTENT

**Supporting Information (DOC)**

**Supplemental Table 1.** Primer pairs used for mutagenesis a wild-type CKB vector.

**Supplemental Table 2.** Reference SNP ID and predicted impact scores (SIFT and PolyPen) of the fifteen naturally mutations investigated.

**Supplemental Figure 1.** Michaelis-Menten plots of the mutant CKB enzymes.

**Supplemental Figure 2.** Melting curves of remaining mutant CKB proteins.

**Supplemental Table 3.** A list of all nucleotide and small molecule kinases that were identified by mass spectrometry-based proteomics in brain and liver.

**Supplemental Table 4.** List of nucleotide and small molecule kinases in brain with differential abundance between sexes and/or genotypes.

**Supplemental Table 5.** List of nucleotide and small molecule kinases in liver with differential abundance between sexes and/or genotypes.

## AUTHOR INFORMATION

### Author Contributions

CDE: conducted experiments, analyzed data, wrote manuscript

EPM: experimental design, wrote manuscript

NNB: experimental design, analyzed data, wrote manuscript

BCO: experimental design, wrote manuscript

### Funding Sources

NIH T32 GM135083

NIH R01 AG064908

## ABBREVIATIONS

CKB: creatine kinase brain-type
TFV: tenofovir
TFV-MP: tenofovir-monophosphate
TFV-DP: tenofovir-diphosphate
WT: wild-type
HIV: human immunodeficiency virus
HBV: hepatitis B virus
PCR: polymerase chain reaction

