## Supplemental for "Tenofovir Activation is Diminished in the Brain and Liver of Creatine Kinase Brain-Type Knockout Mice"

**Supporting Information**

**Supplemental Table 1.** Primer pairs used for mutagenesis a wild-type CKB vector.[1]

| Mutation | Nucleotide Change | Direction | Primer Sequence 5' to 3' |
|---|---|---|---|
| C74S | t220a | Forward | TGACCGTGGGCAGCGTGGCGGGC |
| | | Reverse | CGCCCGCCACGCTGCCCACGGTCA |
| R96P | g287c | Forward | CCCATCATCGAGGACCCGCACGGCGGCTAC |
| | | Reverse | GTAGCCGCCGTGCGGGTCCTCGATGATGGG |
| S128R | a382c | Forward | CTACGTGCTGCGCTCGCGGGTGCGCAC |
| | | Reverse | GTGCGCACCCGCGAGCGCAGCACGTAG |
| R132H | g395a | Forward | CTCGCGGGTGCACACGGGCCGCAG |
| | | Reverse | CTGCGGCCCGTGTGCACCCGCGAG |
| R172G | c514g & a516c | Forward | CGACCTGGCGGGCGGCTACTACGCGC |
| | | Reverse | GCGCGTAGTAGCCGCCCGCCAGGTCG |
| R172P | g515c & a516g | Forward | CGACCTGGCGGGCCCGTACTACGCGCTC |
| | | Reverse | GAGCGCGTAGTACGGGCCCGCCAGGTC |
| R236Q | g707a | Forward | GGAGGACCACCTGCAGGTCATCTCCATGCAGAAGGGG |
| | | Reverse | CCCCCTTCTGCATGGAGATGACCTGCAGGTGGTCCTCC |
| D268H | g802c | Forward | CAAGTCTAAGCACTATGAGTTCATGTGGAACCCTCACCTGGGC |
| | | Reverse | GCCCAGGTGAGGGTTCCACATGAACTCATAGTGCTTAGACTTG |
| C283S | t847a | Forward | TGGGCTACATCCTCACCAGCCCATCCAACCTGGGC |
| | | Reverse | GCCCAGGTTGGATGGGCTGGTGAGGATGTAGCCCA |
| R292Q | g875a | Forward | GGCACCGGGCTGCAGGCAGGTGTGC |
| | | Reverse | GCACACCTGCCTGCAGCCCGGTGCCC |
| H296R | a887g | Forward | GCTGCGGGCAGGTGTGCGTATCAAGCTGCCC |
| | | Reverse | GGGCAGCTTGATACGCACACCTGCCCGCAGCC |
| H296P | a887c & t888g | Forward | GCGGGCAGGTGTGCCGATCAAGCTGCCC |
| | | Reverse | GGGCAGCTTGATCGGCACACCTGCCCGCAGC |
| P300L | c899t & c900g | Forward | GGCAGGTGTGCATATCAAGCTGCTGAACCTGGGCAAGC |
| | | Reverse | GCTTGCCCAGGTTCAGCAGCTTGATATGCACACCTGCCCG |
| T327R | c980g | Forward | GCGGTGTGGACAGGGCTGCGGTGG |
| | | Reverse | CCACCGCAGCCCTGTCCACACCGC |
| I361T | t1082c | Forward | GTGGACGGAGTGAAGCTGCTCACCCGAGATGGAGCAGC |
| | | Reverse | GCTGCTCCATCTCGGTGAGCAGCTTCACTCCGTCCACC |
| E368G | a1103g | Forward | GCAGCGGCTGGGGCAGGGCCAGG |
| | | Reverse | GCCTGGCCCTGCCCCAGCCGCTGC |

[1] Red font in the primer sequences indicates the nucleotide change that will result in the missense mutation

**Supplemental Table 2.** Fifteen naturally occurring missense mutations were selected based on their location in the crystal structure, predicted impact (SIFT and PolyPhen scores), and residue conservation between creatine kinase enzymes.

| Mutation | Location in Crystal Structure | Reference SNP ID | SIFT Prediction | PolyPhen Prediction (PP2HDIV) | Conservation across Creatine Kinase family |
|---|---|---|---|---|---|
| C74S | Creatine binding pocket | rs1350946529 | Damaging (0.005) | Probably Damaging (0.993) | Conserved in CKM, Met in CKMT1/2 |
| R96P | Creatine binding pocket | rs753594980 | Damaging (0.002) | Probably Damaging (0.989) | Conserved |
| S128R | ADP binding pocket | rs747114408 | Damaging (0.0) | Possibly Damaging (0.936) | Conserved |
| R132H | Active site arginine | rs1467790706 | Damaging (0.019) | Possibly Damaging (0.467) | Conserved |
| R172G | Dimer interface | rs745508721 | Damaging (0.002) | unknown | Conserved in CKB and CKMT1/2, Lys in CKM |
| R172P | Dimer interface | rs1389333856 | Damaging (0.002) | Possibly Damaging (0.599) | |
| R236Q | Active site arginine | rs1489604594 | Damaging (0.003) | Possibly Damaging (0.948) | Conserved |
| D268H | Dimer interface | rs770230629 | Damaging (0.001) | Benign (0.052) | Gly in CKM and CKMT1/2 |
| R292Q | Active site arginine | rs1367035745 | Damaging (0.0) | Probably Damaging (0.963) | Conserved |
| H296R | ADP binding pocket | rs758039698 | Damaging (0.001) | Probably Damaging (0.001) | Conserved |
| H296P | ADP binding pocket | rs758039698 | Damaging (0.001) | Probably Damaging (1.0) | |
| P300L | External surface | rs764629320 | Damaging (0.001) | Possibly Damaging (0.477) | Conserved in CKB and CKMT1/2, Ala in CKM |
| T327R | Dimer interface | rs769802932 | Damaging (0.0) | Probably Damaging (1.0) | Conserved |
| I361T | External surface | rs145462676 | Damaging (0.009) | Probably Damaging (0.974) | Conserved in CKB and CKMT1, Val in CKM and CKMT2 |
| E368G | External surface | rs748950650 | Damaging (0.007) | Possibly Damaging (0.697) | Conserved |

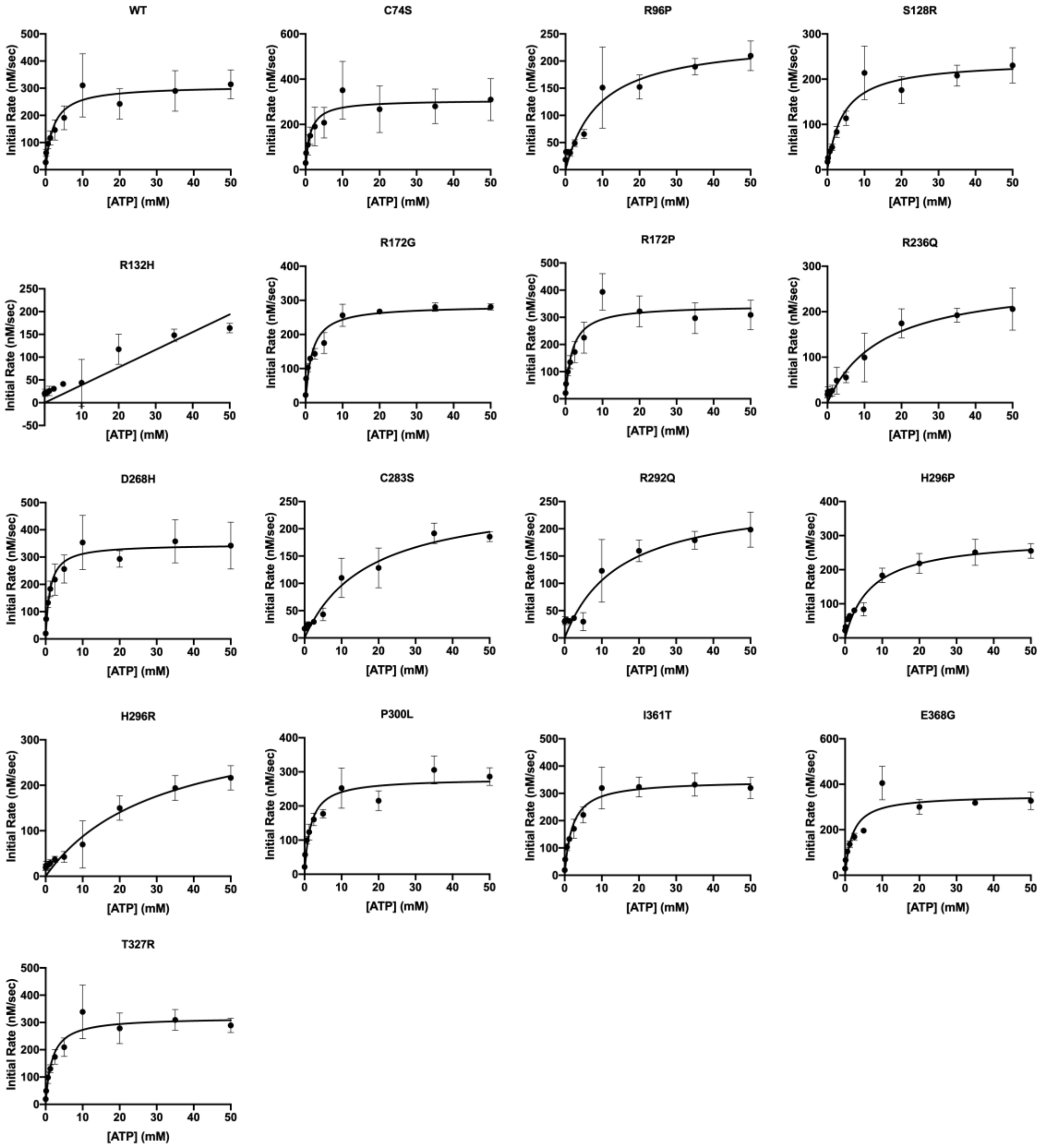


**Supplemental Figure 1.** Recombinantly expressed mutant CKB enzymes were coupled in a pyruvate kinase/lactate dehydrogenase enzyme system to obtain Michaelis-Menten kinetic parameters. To obtain Michaelis-Menten plots, initial rates at varying ATP concentrations was determined and plotted against the concentration of ATP. Mutant CKB assays were performed with n = 3, while WT CKB was performed n = 8. Data obtained from R132H better fit a linear regression than the Michaelis-Menten equation.

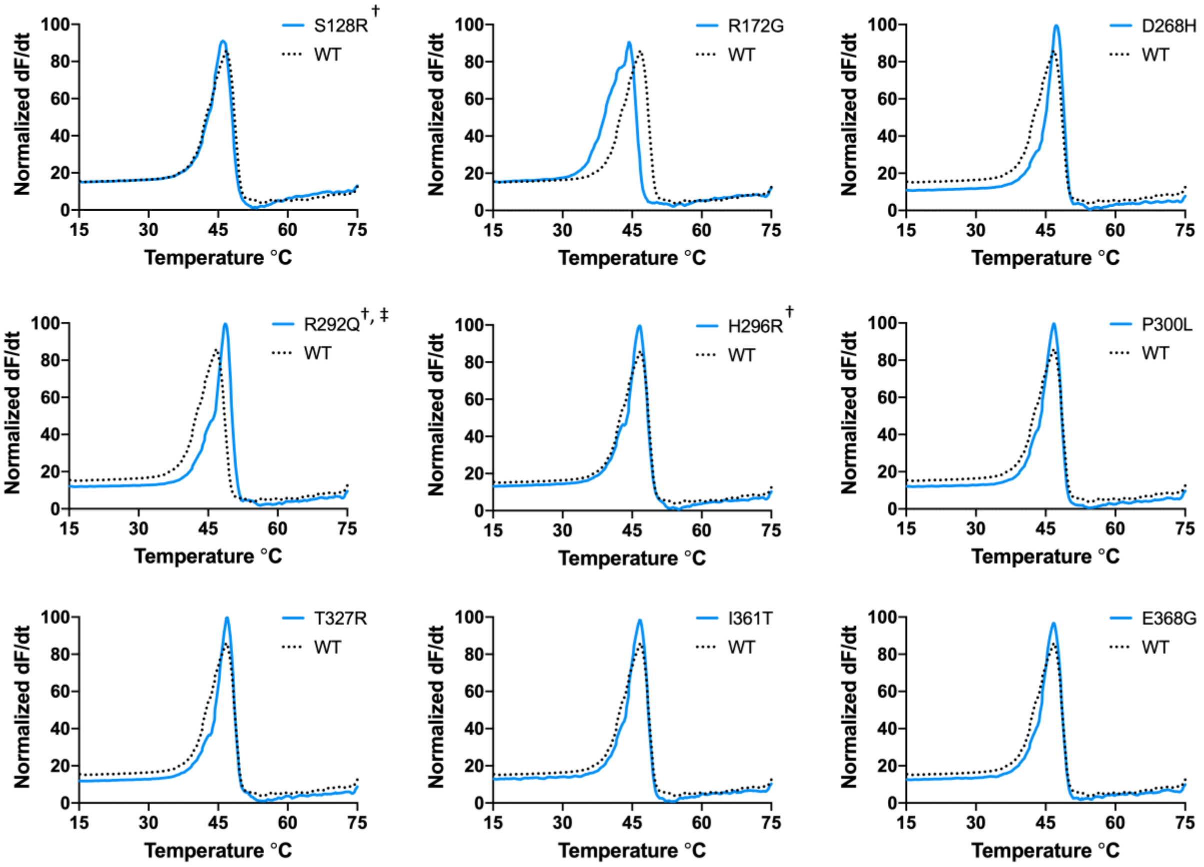


**Supplemental Figure 2.** Differential scanning fluorimetry was utilized to obtain the melting curves of mutant CKB proteins. The melting curves of S128R, R172G, D268H, R292Q, H296R, P300L, T327R, I361T, and E368G do not contain prominent double peaks or shouldering with a statistically different melting temperature when compared to the wild-type curve.

**Supplemental Table 3.** A list of all nucleotide and small molecule kinases that were identified by mass spectrometry-based proteomics in brain and liver.[2]

| UniProt Accession | Gene Symbol | Protein Name | Tissue |
|---|---|---|---|
| P55264 | Adk | Adenosine kinase | Brain, Liver |
| Q8VDL4 | Adpgk | ADP-dependent glucokinase | Brain, Liver |
| Q9ESW4 | Agk | Acylglycerol kinase, mitochondrial | Brain, Liver |
| Q9R0Y5 | Ak1 | Adenylate kinase isoenzyme 1 | Brain |
| Q9WTP6 | Ak2 | Adenylate kinase 2, mitochondrial | Brain, Liver |
| Q9WTP7 | Ak3 | GTP:AMP phosphotransferase AK3, mitochondrial | Brain, Liver |
| Q9WUR9 | Ak4 | Adenylate kinase 4, mitochondrial | Brain, Liver |
| Q920P5 | Ak5 | Adenylate kinase isoenzyme 5 | Brain |
| P08030 | Aprt | Adenine phosphoribosyltransferase | Brain |
| Q9Z0S1 | Bpnt1 | 3'(2'),5'-bisphosphate nucleotidase 1 | Brain, Liver |
| Q8VCF1 | Cant1 | Soluble calcium-activated nucleotidase 1 | Liver |
| O54804 | Chka | Choline kinase alpha | Brain, Liver |
| O55229 | Chkb | Choline/ethanolamine kinase | Brain, Liver |
| Q04447 | Ckb | Creatine kinase B-type | Brain, Liver |
| P07310 | Ckm | Creatine kinase M-type | Brain |
| P30275 | Ckmt1 | Creatine kinase U-type, mitochondrial | Brain |
| Q9DBP5 | Cmpk1 | UMP-CMP kinase | Brain, Liver |
| Q3U5Q7 | Cmpk2 | UMP-CMP kinase 2, mitochondrial | Brain, Liver |
| P16330 | Cnp | 2',3'-cyclic-nucleotide 3'-phosphodiesterase | Brain, Liver |
| Q60936 | Coq8a | Atypical kinase COQ8A, mitochondrial | Brain, Liver |
| Q566J8 | Coq8b | Atypical kinase COQ8B, mitochondrial | Liver |
| Q8BHC4 | Dcakd | Dephospho-CoA kinase domain-containing protein | Brain, Liver |
| P43346 | Dck | Deoxycytidine kinase | Brain |
| Q6NS52 | Dgkb | Diacylglycerol kinase beta | Brain |
| E9PUQ8 | Dgkd | Diacylglycerol kinase delta | Brain |
| Q9R1C6 | Dgke | Diacylglycerol kinase epsilon | Brain |
| Q91WG7 | Dgkg | Diacylglycerol kinase gamma | Brain |
| D3YXJ0 | Dgkh | Diacylglycerol kinase eta | Brain |
| D3YWQ0 | Dgki | Diacylglycerol kinase iota | Brain |
| Q6P5E8 | Dgkq | Diacylglycerol kinase theta | Brain |
| Q80UP3 | Dgkz | Diacylglycerol kinase zeta | Brain |
| Q9QX60 | Dguok | Deoxyguanosine kinase, mitochondrial | Brain, Liver |
| Q80VJ3 | Dnph1 | 2'-deoxynucleoside 5'-phosphate N-hydrolase 1 | Liver |
| P97930 | Dtymk | Thymidylate kinase | Brain, Liver |
| Q9CQ43 | Dut | Deoxyuridine 5'-triphosphate nucleotidohydrolase | Liver |
| P06802 | Enpp1 | Ectonucleotide pyrophosphatase/phosphodiesterase family member 1 | Brain, Liver |
| Q6DYE8 | Enpp3 | Ectonucleotide pyrophosphatase/phosphodiesterase family member 3 | Liver |
| P55772 | Entpd1 | Ectonucleoside triphosphate diphosphohydrolase 1 | Brain |
| O55026 | Entpd2 | Ectonucleoside triphosphate diphosphohydrolase 2 | Brain |
| Q8BFW6 | Entpd3 | Ectonucleoside triphosphate diphosphohydrolase 3 | Brain |

[2] There are 50 uniquely found in brain, 27 unique to liver, and 73 that are found in both tissues.

| | | | |
|---|---|---|---|
| Q9WUZ9 | Entpd5 | Ectonucleoside triphosphate diphosphohydrolase 5 | Liver |
| Q3U0P5 | Entpd6 | Ectonucleoside triphosphate diphosphohydrolase 6 | Brain |
| Q9D4V0 | Etnk1 | Ethanolamine kinase 1 | Brain |
| A7MCT6 | Etnk2 | Ethanolamine kinase 2 | Liver |
| Q7TMC8 | Fcsk | L-fucose kinase | Brain, Liver |
| A2AJL3 | Fggy | FGGY carbohydrate kinase domain-containing protein | Liver |
| Q9R0N0 | Galk1 | Galactokinase | Brain, Liver |
| Q68FH4 | Galk2 | N-acetylgalactosamine kinase | Brain, Liver |
| P52792 | Gck | Hexokinase-4 | Liver |
| Q64516 | Gk | Glycerol kinase | Brain, Liver |
| Q8QZY2 | Glyctk | Glycerate kinase | Liver |
| Q91WG8 | Gne | Bifunctional UDP-N-acetylglucosamine 2-epimerase/N-acetylmannosamine kinase | Brain, Liver |
| Q64520 | Guk1 | Guanylate kinase | Brain, Liver |
| Q3SXD3 | Hddc2 | 5'-deoxynucleotidase HDDC2 | Brain |
| P70349 | Hint1 | Adenosine 5'-monophosphoramidase HINT1 | Brain |
| Q9D0S9 | Hint2 | Adenosine 5'-monophosphoramidase HINT2 | Brain |
| Q9CPS6 | Hint3 | Adenosine 5'-monophosphoramidase HINT3 | Brain |
| P17710 | Hk1 | Hexokinase-1 | Brain |
| O08528 | Hk2 | Hexokinase-2 | Brain |
| Q3TRM8 | Hk3 | Hexokinase-3 | Liver |
| P00493 | Hprt1 | Hypoxanthine-guanine phosphoribosyltransferase | Brain |
| Q8R0J8 | Idnk | Probable gluconokinase | Liver |
| Q6PD10 | Ip6k1 | Inositol hexakisphosphate kinase 1 | Brain |
| Q8BYN3 | Itpk1 | Inositol-tetrakisphosphate 1-kinase | Brain |
| Q8R071 | Itpka | Inositol-trisphosphate 3-kinase A | Brain |
| Q7TS72 | Itpkc | Inositol-trisphosphate 3-kinase C | Brain |
| P97328 | Khk | Ketohexokinase | Brain, Liver |
| Q6RHR9 | Magi1 | Membrane-associated guanylate kinase, WW and PDZ domain-containing protein 1 | Brain |
| Q99JF5 | Mvd | Diphosphomevalonate decarboxylase | Brain, Liver |
| Q9R008 | Mvk | Mevalonate kinase | Brain, Liver |
| P58058 | Nadk | NAD kinase | Brain, Liver |
| Q8C5H8 | Nadk2 | NAD kinase 2, mitochondrial | Brain, Liver |
| Q9QZ08 | Nagk | N-acetyl-D-glucosamine kinase | Brain, Liver |
| Q8CC86 | Naprt | Nicotinate phosphoribosyltransferase | Brain, Liver |
| P15532 | Nme1 | Nucleoside diphosphate kinase A | Brain, Liver |
| Q01768 | Nme2 | Nucleoside diphosphate kinase B | Brain, Liver |
| Q9WV85 | Nme3 | Nucleoside diphosphate kinase 3 | Brain, Liver |
| Q9EPA7 | Nmnat1 | Nicotinamide/nicotinic acid mononucleotide adenylyltransferase 1 | Liver |
| Q99JR6 | Nmnat3 | Nicotinamide/nicotinic acid mononucleotide adenylyltransferase 3 | Brain, Liver |
| Q91W63 | Nmrk1 | Nicotinamide riboside kinase 1 | Liver |
| Q9JM14 | Nt5c | 5'(3')-deoxyribonucleotidase, cytosolic type | Brain, Liver |
| A3KFX0 | Nt5c1a | Cytosolic 5'-nucleotidase 1A | Brain |
| Q3V1L4 | Nt5c2 | Cytosolic purine 5'-nucleotidase | Brain, Liver |
| Q9D020 | Nt5c3a | Cytosolic 5'-nucleotidase 3A | Brain, Liver |
| Q3UFY7 | Nt5c3b | 7-methylguanosine phosphate-specific 5'-nucleotidase | Brain |
| Q8C5P5 | Nt5dc1 | 5'-nucleotidase domain-containing protein 1 | Liver |
| Q3UHB1 | Nt5dc3 | 5'-nucleotidase domain-containing protein 3 | Brain |

| | | | |
|---|---|---|---|
| Q61503 | Nt5e | 5'-nucleotidase | Brain, Liver |
| Q8VCE6 | Nt5m | 5'(3')-deoxyribonucleotidase, mitochondrial | Brain |
| Q9CQA9 | Ntpcr | Cancer-related nucleoside-triphosphatase homolog | Liver |
| Q8BG93 | Nudt15 | Nucleotide triphosphate diphosphatase NUDT15 | Brain |
| Q9CWD3 | Nudt17 | Nucleoside diphosphate-linked moiety X motif 17 | Brain |
| P56380 | Nudt2 | Bis(5'-nucleosyl)-tetraphosphatase [asymmetrical] | Brain, Liver |
| Q9DCL9 | Paics | Bifunctional phosphoribosylaminoimidazole carboxylase/phosphoribosylaminoimidazole succinocarboxamide synthetase | Brain, Liver |
| Q8K4K6 | Pank1 | Pantothenate kinase 1 | Brain, Liver |
| Q7M753 | Pank2 | Pantothenate kinase 2, mitochondrial | Brain |
| Q8R2W9 | Pank3 | Pantothenate kinase 3 | Liver |
| Q60967 | Papss1 | Bifunctional 3'-phosphoadenosine 5'-phosphosulfate synthase 1 | Brain, Liver |
| O88428 | Papss2 | Bifunctional 3'-phosphoadenosine 5'-phosphosulfate synthase 2 | Brain, Liver |
| Q8BH04 | Pck2 | Phosphoenolpyruvate carboxykinase [GTP], mitochondrial | Brain, Liver |
| Q61481 | Pde1a | Dual specificity calcium/calmodulin-dependent 3',5'-cyclic nucleotide phosphodiesterase 1A | Brain |
| Q01065 | Pde1b | Dual specificity calcium/calmodulin-dependent 3',5'-cyclic nucleotide phosphodiesterase 1B | Brain |
| Q64338 | Pde1c | Dual specificity calcium/calmodulin-dependent 3',5'-cyclic nucleotide phosphodiesterase 1C | Brain |
| Q8K183 | Pdxk | Pyridoxal kinase | Brain, Liver |
| Q5SUR0 | Pfas | Phosphoribosylformylglycinamidine synthase | Liver |
| P70266 | Pfkfb1 | 6-phosphofructo-2-kinase/fructose-2,6-bisphosphatase 1 | Liver |
| P70265 | Pfkfb2 | 6-phosphofructo-2-kinase/fructose-2,6-bisphosphatase 2 | Brain, Liver |
| P12382 | Pfkl | ATP-dependent 6-phosphofructokinase, liver type | Brain, Liver |
| P47857 | Pfkm | ATP-dependent 6-phosphofructokinase, muscle type | Brain, Liver |
| Q9WUA3 | Pfkp | ATP-dependent 6-phosphofructokinase, platelet type | Brain, Liver |
| P09411 | Pgk1 | Phosphoglycerate kinase 1 | Brain, Liver |
| Q2TBE6 | Pi4k2a | Phosphatidylinositol 4-kinase type 2-alpha | Brain, Liver |
| E9Q3L2 | Pi4ka | Phosphatidylinositol 4-kinase alpha | Brain, Liver |
| Q8BKC8 | Pi4kb | Phosphatidylinositol 4-kinase beta | Brain, Liver |
| Q61194 | Pik3c2a | Phosphatidylinositol 4-phosphate 3-kinase C2 domain-containing subunit alpha | Brain, Liver |
| Q6PF93 | Pik3c3 | Phosphatidylinositol 3-kinase catalytic subunit type 3 | Brain, Liver |
| P42337 | Pik3ca | Phosphatidylinositol 4,5-bisphosphate 3-kinase catalytic subunit alpha isoform | Brain |
| Q8BTI9 | Pik3cb | Phosphatidylinositol 4,5-bisphosphate 3-kinase catalytic subunit beta isoform | Brain |
| O70172 | Pip4k2a | Phosphatidylinositol 5-phosphate 4-kinase type-2 alpha | Brain, Liver |
| Q80XI4 | Pip4k2b | Phosphatidylinositol 5-phosphate 4-kinase type-2 beta | Brain, Liver |
| Q91XU3 | Pip4k2c | Phosphatidylinositol 5-phosphate 4-kinase type-2 gamma | Brain, Liver |
| P70182 | Pip5k1a | Phosphatidylinositol 4-phosphate 5-kinase type-1 alpha | Brain |
| O70161 | Pip5k1c | Phosphatidylinositol 4-phosphate 5-kinase type-1 gamma | Brain |
| P53657 | Pklr | Pyruvate kinase PKLR | Liver |
| P52480 | Pkm | Pyruvate kinase PKM | Brain, Liver |
| Q9D1G2 | Pmvk | Phosphomevalonate kinase | Liver |
| P23492 | Pnp | Purine nucleoside phosphorylase | Brain, Liver |
| Q8VDG5 | Ppcs | Phosphopantothenate--cysteine ligase | Liver |
| A2ARP1 | Ppip5k1 | Inositol hexakisphosphate and diphosphoinositol-pentakisphosphate kinase 1 | Brain |
| Q6ZQB6 | Ppip5k2 | Inositol hexakisphosphate and diphosphoinositol-pentakisphosphate kinase 2 | Liver |

| | | | |
|---|---|---|---|
| Q9D7G0 | Prps1 | Ribose-phosphate pyrophosphokinase 1 | Brain, Liver |
| Q9CS42 | Prps2 | Ribose-phosphate pyrophosphokinase 2 | Brain, Liver |
| Q8R1Q9 | Rbks | Ribokinase | Liver |
| Q8CFV9 | Rfk | Riboflavin kinase | Liver |
| P07742 | Rrm1 | Ribonucleoside-diphosphate reductase large subunit | Brain |
| Q6PEE3 | Rrm2b | Ribonucleoside-diphosphate reductase subunit M2 B | Brain |
| Q60710 | Samhd1 | Deoxynucleoside triphosphate triphosphohydrolase SAMHD1 | Brain, Liver |
| Q8BH69 | Sephs1 | Selenide, water dikinase 1 | Brain, Liver |
| P97364 | Sephs2 | Selenide, water dikinase 2 | Brain, Liver |
| Q9D5J6 | Shpk | Sedoheptulokinase | Liver |
| Q9JIM1 | Slc29a1 | Equilibrative nucleoside transporter 1 | Brain, Liver |
| Q61672 | Slc29a2 | Equilibrative nucleoside transporter 2 | Brain |
| P70158 | Smpdl3a | Acid sphingomyelinase-like phosphodiesterase 3a | Brain, Liver |
| Q9JIA7 | Sphk2 | Sphingosine kinase 2 | Brain, Liver |
| Q9R088 | Tk2 | Thymidine kinase 2, mitochondrial | Brain, Liver |
| Q8VC30 | Tkfc | Triokinase/FMN cyclase | Brain, Liver |
| Q9R0M5 | Tpk1 | Thiamin pyrophosphokinase 1 | Brain, Liver |
| P52623 | Uck1 | Uridine-cytidine kinase 1 | Brain, Liver |
| Q91YL3 | Uckl1 | Uridine-cytidine kinase-like 1 | Brain |
| Q3TNA1 | Xylb | Xylulose kinase | Liver |

**Supplemental Table 4.** List of nucleotide and small molecule kinases in brain with differential abundance between sexes and/or genotypes.[3]

| WT Male vs. KO Male | | WT Female vs. KO Female | | WT Male vs. WT Female | | KO Male vs. KO Female |
|---|---|---|---|---|---|---|
| Ak2 | **Paics** | **Ak1** | **Mvd** | Ak1 | **Pfkm** | **Ak2** |
| Adpgk | Pank1 | Adpgk | Mvk | Ak3 | **Pfkp** | Ak3 |
| **Ak3** | Papss1 | Ak2 | Nadk | Ak4 | Pgk1 | **Ckmt1** |
| **Chkb** | **Papss2** | **Ak3** | Naprt | **Chka** | **Pkm** | Cmpk2 |
| Ckb | Pdxk | **Ak4** | Nme1 | **Chkb** | **Prps1** | **Dguok** |
| Ckm | Pfkl | **Ak5** | Nme2 | Ckb | **Prps2** | Enpp1 |
| **Cmpk2** | Pfkp | Chka | **Papss2** | **Dguok** | **Sephs1** | Entpd2 |
| Dguok | Pgk1 | Chkb | Pfkl | **Enpp1** | **Smpdl3a** | **Gk** |
| **Dtymk** | Pkm | Ckb | Pfkm | **Entpd1** | | Itpka |
| **Entpd1** | Ppip5k1 | Ckm | Pfkp | **Entpd3** | | Nadk |
| Entpd2 | Prps2 | **Ckmt1** | **Pgk1** | **Etnk1** | | **Nagk** |
| Gk | Tpk1 | Dguok | Pik3ca | Galk1 | | **Nme1** |
| Guk1 | **Uckl1** | **Dtymk** | Pkm | Guk1 | | **Nme2** |
| **Hk1** | | Enpp1 | Prps1 | Hk1 | | Nme3 |
| **Ip6k1** | | Entpd2 | Prps2 | Itpka | | Nudt17 |
| Khk | | Entpd3 | Sephs1 | Mvd | | Papss2 |
| Mvk | | Etnk1 | Sephs2 | **Nme1** | | **Pdxk** |
| Nme1 | | Fcsk | Smpdl3a | **Nme2** | | **Pgk1** |
| Nme2 | | Gk | Tk2 | Nudt15 | | **Pkm** |
| **Nme3** | | **Hk1** | Tpk1 | Papss2 | | **Prps1** |
| Nudt15 | | Khk | **Uckl1** | **Pfkl** | | **Prps2** |

[3] Listed are Gene IDs of proteins that use ATP as a substrate to phosphorylate nucleotides or small molecules, or chemically modify ATP. Additionally, these proteins have a reaction compatible with the structure of TFV metabolites. Comparisons examine changes between sexes and genotypes (n = 7 each group). Statistical analyses were conducted using SimpliFi. All proteins have q-value < 0.05. Bold font indicates the protein in higher in abundance in the first experimental group.

**Supplemental Table 5.** List of nucleotide and small molecule kinases in liver with differential abundance between sexes and/or genotypes.[4]

| WT Male vs. KO Male | | WT Female vs. KO Female | | WT Male vs. WT Female | | KO Male vs. KO Female |
|---|---|---|---|---|---|---|
| Ak2 | **Pgk1** | **Adk** | Nme2 | Adk | Pklr | Ak2 |
| **Ak3** | Pklr | Ak2 | **Nmnat1** | Ak3 | **Pkm** | Cant1 |
| **Ak4** | **Ppip5k2** | **Ak3** | **Nmnat3** | Ak4 | Pmvk | Cmpk1 |
| **Cant1** | Prps2 | **Ak4** | **Nmrk1** | Ckb | **Prps1** | **Cmpk2** |
| Ckb | Rbks | **Chkb** | **Pank1** | Cmpk1 | **Sephs2** | **Enpp1** |
| **Cmpk1** | **Rfk** | **Cmpk1** | **Papss2** | **Dnph1** | Tkfc | Enpp3 |
| Dguok | Sephs1 | Dguok | Pdxk | **Enpp1** | **Tpk1** | Entpd5 |
| **Enpp3** | Sephs2 | Dnph1 | **Pfas** | Enpp3 | Xylb | **Fggy** |
| **Entpd5** | **Tkfc** | **Dtymk** | **Pfkfb1** | Entpd5 | | **Gk** |
| Fggy | Tpk1 | **Entpd5** | Pfkfb2 | **Fggy** | | Idnk |
| **Galk2** | **Xylb** | Fcsk | Pfkl | Galk1 | | **Mvd** |
| **Gck** | | Fggy | **Pgk1** | Galk2 | | **Mvk** |
| Gk | | **Galk1** | **Pklr** | Gck | | **Nadk** |
| **Glyctk** | | **Galk2** | Pkm | **Gk** | | Paics |
| **Idnk** | | **Gck** | **Pmvk** | Glyctk | | **Papss2** |
| Khk | | Gk | **Ppip5k2** | Guk1 | | Pdxk |
| **Mvd** | | **Glyctk** | Prps2 | Hk3 | | Pfkfb1 |
| Nadk | | **Guk1** | Rbks | Idnk | | **Pfkl** |
| Nme2 | | **Hk3** | **Rfk** | **Khk** | | **Pklr** |
| **Nmnat1** | | **Idnk** | Sephs1 | **Mvk** | | **Pkm** |
| **Nmnat3** | | Khk | Sephs2 | **Nadk** | | Ppip5k2 |
| **Nmrk1** | | **Mvd** | **Shpk** | Papss1 | | **Prps1** |
| **Pank1** | | Nadk | **Tkfc** | Pfas | | **Prps2** |
| **Papss2** | | **Nadk2** | Tpk1 | Pfkfb1 | | Tkfc |
| Pfkfb2 | | Naprt | **Uck1** | **Pfkl** | | |
| Pfkl | | Nme1 | **Xylb** | Pgk1 | | |

[4] Listed are Gene IDs of proteins that use ATP as a substrate to phosphorylate nucleotides or small molecules, or chemically modify ATP. Additionally, these proteins have a reaction compatible with the structure of TFV metabolites. Comparisons examine changes between sexes and genotypes (n = 7 each group). Statistical analyses were conducted using SimpliFi. All proteins have q-value < 0.05. Bold font indicates the protein in higher in abundance in the first experimental group.
